# Appropriate tension sensitivity of α-catenin ensures rounding morphogenesis of epithelial spheroids

**DOI:** 10.1101/2021.10.27.466008

**Authors:** Ryosuke Nishimura, Kagayaki Kato, Misako Saida, Yasuhiro Kamei, Masahiro Takeda, Hiromi Miyoshi, Yutaka Yamagata, Yu Amano, Shigenobu Yonemura

**Affiliations:** Department of Cell Biology, Graduate School of Medical Sciences, Tokushima University, 3-18-15 Kuramoto-cho, Tokushima, Tokushima 770-8503, Japan; Exploratory Research Center on Life and Living Systems (ExCELLS), National Institutes of Natural Sciences, 38 Nishigonaka, Myodaiji, Okazaki, Aichi 444-8585, Japan; Spectrography and Bioimaging Facility, National Institute for Basic Biology, 38 Nishigonaka, Myodaiji, Okazaki, Aichi 444-8585, Japan; Department of Basic Biology, School of Life Science, The Graduate University for Advanced Studies (SOKENDAI), 38, Nishigonaka, Myodaiji, Okazaki, Japan; Ultra High Precision Optics Technology Team/Advanced Manufacturing Support Team, RIKEN, 2-1 Hirosawa, Wako, Saitama 351-0198, Japan; Center for Advance Photonics, RIKEN, 2-1 Hirosawa, Wako, Saitama 351-0198, Japan; Applied Mechanobiology Laboratory, Faculty of Systems Design, Tokyo Metropolitan University, 1-1 Minami-Osawa, Hachioji, Tokyo,192-0397, Japan; Department of Bioscience, Kwansei Gakuin University, Sanda, Hyogo 669-1337, Japan; Ultrastructural Research Team, RIKEN Center for Biosystems Dynamics Research, Kobe, Hyogo, Japan

**Keywords:** α-catenin, vinculin, adherens junction, epithelial morphogenesis, mechanotransduction

## Abstract

The adherens junction (AJ) is an actin filament-anchoring junction. It plays a central role in epithelial morphogenesis through cadherin-based recognition and adhesion among cells. The stability and plasticity of AJs are required for the morphogenesis. An actin-binding α-catenin is an essential component of the cadherin-catenin complex and functions as a tension transducer that changes its conformation and induces AJ development in response to tension. Despite much progress in understanding molecular mechanisms of tension sensitivity of α-catenin, its significance on epithelial morphogenesis is still unknown. Here we show that the tension sensitivity of α-catenin is essential for epithelial cells to form round spheroids through proper multicellular rearrangement. Using a novel *in vitro* suspension culture model, we found that epithelial cells form round spheroids even from rectangular-shaped cell masses with high aspect ratios without using high tension and that hypersensitive mutants affected this morphogenesis. Analyses of AJ formation and cellular tracking during rounding morphogenesis showed cellular rearrangement, probably through AJ remodeling. The rearrangement occurs at the cell mass level, but not single-cell level. Hypersensitive α-catenin mutant-expressing cells did not show cellular rearrangement at the cell mass level, suggesting that proper AJ plasticity requires appropriate tension sensitivity of α-catenin.

## Introduction

The morphogenesis mechanism in multicellular organisms is one of the most important research topics for developmental and cell biologists. The epithelial tissue that separates the inside and outside of the living environment contributes to a significant part of morphogenesis. It is a sheet-like structure composed of epithelial cells connected by several types of cell-cell junctions. The epithelial sheet can stretch, bend, or partly protrude to form basic structures common among species, such as grooves, hollow structures, and branching (1). Since the early 20th century, the morphogenesis of multicellular organisms has been studied. Experiments on the separation and reassembly of sea urchin embryos or sponges indicated that cells can sort themselves depending on tissue specificity and spontaneously reassemble their tissues (2, 3). Such features were also observed in vertebrates, and differences in cell-cell adhesive force (4) or surface tension (5) of cells were proposed as the primary factor causing sorting in mammalian cells. Subsequent studies showed that the adhesive force increases in proportion to the number of adhesion molecules and inverse proportion to cortical tension (which decreases adhesive area), and differences in the total adhesive force causes sorting (6, 7). The actual morphogenesis in living organisms is more complex, and many studies have been conducted using various organisms. They revealed the importance of spatiotemporal regulation of programmed cell death (8), cell division (9), cell migration (10), and cell deformation (11) for morphogenesis. Cell division and apoptosis contribute to local cell number and rearrangement regulation, while cell migration contributes to tissue rearrangement on a larger scale. Signaling plays an important role in determining the timing of these changes through communication between distant cells (12).

Many cellular deformations that directly contribute to large deformations of epithelial tissues depend on the forces generated by actomyosin in individual cells (13, 14). Recently, the involvement of force in epithelial morphogenesis has been intensively studied using theoretical approaches. Many mathematical models have been established to predict morphological changes in epithelial tissues in response to mechanical stimuli (15–19). Because precision manipulation or measurement of force *in vivo* is highly challenging, theoretical modeling occupies an essential position in understanding multicellular behavior during morphogenesis. These experimental and theoretical investigations gave a better understanding of the generation and transmission of tension and its responses to multicellular organisms’ morphogenesis.

Adherens junctions (AJs) are initially formed when cells come into contact with each other and serve as sites for transmitting tension between cells. AJs are cadherin-based and formed near the apical region of cell boundaries, where the actin cytoskeleton is anchored by α-catenin (20). AJs play a vital role in transmitting forces required for cell deformation to the plasma membrane as actin-anchoring sites and transmitting them to adjacent cells (21, 22). The contraction of AJ-anchored actomyosin causes apical constriction at the AJ location. AJs withstand tension and maintain connections between adjacent cells (23). Then, the deformation of individual cells causes the deformation of the entire epithelial sheet. The disruption of either intracellular AJ-actin filament association or extracellular cadherin-cadherin association at AJs causes morphogenetic failure. In the process of epithelial morphogenesis, such as the ventral furrow formation in *Drosophila* embryos and the neural tube in vertebrates, AJs must maintain sufficient structural stability (24).

Many molecules are involved in ensuring the structural stability of AJs. Cadherins enable adhesion to neighboring cells through homophilic binding in their extracellular region, and α-catenin binds to the cytoplasmic region through β-catenin to form the cadherin-catenin complex (25, 26). α-Catenin binds to actin filaments either directly (27, 28) or by actin-binding proteins, including vinculin (29-31). α-Catenin is essential in AJ formation since the loss of α-catenin, or its actin-binding ability, causes failure of AJ formation and loss of actin filament’s association with the complex (32-36). In addition to α-catenin, other actin-binding proteins, such as afadin (37, 38) or ZO-1 (39) accumulate in AJs, and actin filaments are also frequently associated with AJ in tissues. In a typical polarized epithelial cell, the AJ surrounds the apical region of the cell as a belt, and an actin filament bundle develops with it. Many of the above actin-binding proteins can bind to α-catenin. Among them; vinculin binds to α-catenin in a tension-dependent manner (29-31). The importance of vinculin in AJ-associated morphogenesis is evident; loss of vinculin leads to failure of the neural tube closure, a process involving apical constriction described above, and severely affects heart development (40), in which AJs are constantly subjected to high tension.

Under tension-free conditions, α-catenin does not bind to vinculin due to the fact that the α-helix bundle containing the specific α-helix that binds to vinculin is stabilized by intramolecular interactions and cannot interact with vinculin (41, 42). When tension larger than 5 pN is applied to the molecule, the α-helix bundle becomes unstable, allowing the vinculin-binding α-helix to be exposed and bind to vinculin (41, 43). It has been directly confirmed by AFM stretching and real-time TIRF observation that vinculin is recruited when tension is applied to the α-catenin molecule (44). Thus, α-catenin is a force-sensitive vinculin-binding protein. Association/dissociation of actin-binding protein vinculin to α-catenin would contribute to tension-dependent regulation of actin filament-AJ anchoring.

What is the physiological significance of the tension sensitivity of α-catenin? Amino acid residues important for vinculin binding and tension-dependent conformational changes have been identified based on crystal structures, and tension sensitivity mutants have been generated based on this information (41, 45, 46). Several groups have investigated the collective migration of epithelial cells using some of those mutants(37, 47). Those reports show that when tension sensitivity is elevated, allowing the constitutive binding of vinculin to α-catenin, the mutation reduces the speed of collective cell migration. Collective cell migration and local rearrangement between cells are essential for convergent extension (CE) (48). Observations using an α-catenin mutant lacking the vinculin-binding site in zebrafish embryos indicate that viculin’s inability to bind to α-catenin also decreases the rate of collective movement in CE and prevents the completion of CE (49). Thus, although the contexts differ, there is accumulating evidence on the importance of proper binding and dissociation of actin-binding proteins to α-catenin in the planar movement in cells in the same direction. However, the significance of this property of α-catenin in three-dimensional (3D) morphogenesis is almost unknown. Additionally, it is unclear how AJ remodeling is involved in the morphogenesis.

In this study, we used a newly developed morphogenetic model system to investigate the significance of the tension sensitivity of α-catenin on 3D morphogenesis of epithelial cells. We performed analyses at the molecular complex, cellular, and multicellular levels. They suggested that α-catenin’s ability to dissociate actin-binding proteins, in a tension-dependent manner is essential for the normal AJ maturation and the spheroid rounding morphogenesis.

## Results

### Importance of tension sensitivity of α-catenin on 3D morphogenesis of epithelial cells

To clarify the significance of the tension sensitivity of α-catenin, an important component of AJ, in three-dimensional (3D) epithelial morphogenesis, α-catenin mutants with artificially manipulated tension sensitivity for vinculin binding in the middle of the molecule were used in this study (**Fig. 1A**) (29, 41, 46). α-Catenin has a force-sensing device in the middle region, where three bundles consisting of four α-helices are stabilized by interacting with each other (41). The actual mechanistic understanding of the tension sensitivity of α-catenin is as follows: when the molecule is stretched, one of three bundles destabilizes and exposes a vinculin-binding α-helix to enable binding to vinculin. The R326E, R548E, or R551E mutation destabilizes the bundle by weakening the interaction between bundles, and the M319G or L378P mutation constitutively exposes the vinculin-binding α-helix. R2/7 cells, an α-catenin-deficient subclone of DLD-1 cells derived from human colorectal cancer was used (36, 50), expressing each α-catenin mutant stably. These mutants were used previously to investigate the significance of tension sensitivity on the epithelial cell sheet formation (29, 41). It was found that the C-terminus deletion or point mutations in α-catenin, which induces hypersensitivity (**Fig. 1A**), allowed the formation of the tight junction network similar to wild-type (WT), indicating that the essential α-catenin function remains in all mutants. Under conventional two-dimensional (2D) culture conditions, only slight differences were observed in the cell height uniformity and functional cell adhesion timing after cell attachment. Therefore, 3D culture model was used in this study to eliminate the effect of the stiff substrate on the epithelial sheets, and to see morphogenesis that more strongly reflects the nature of the AJ.

**Fig. 1.**
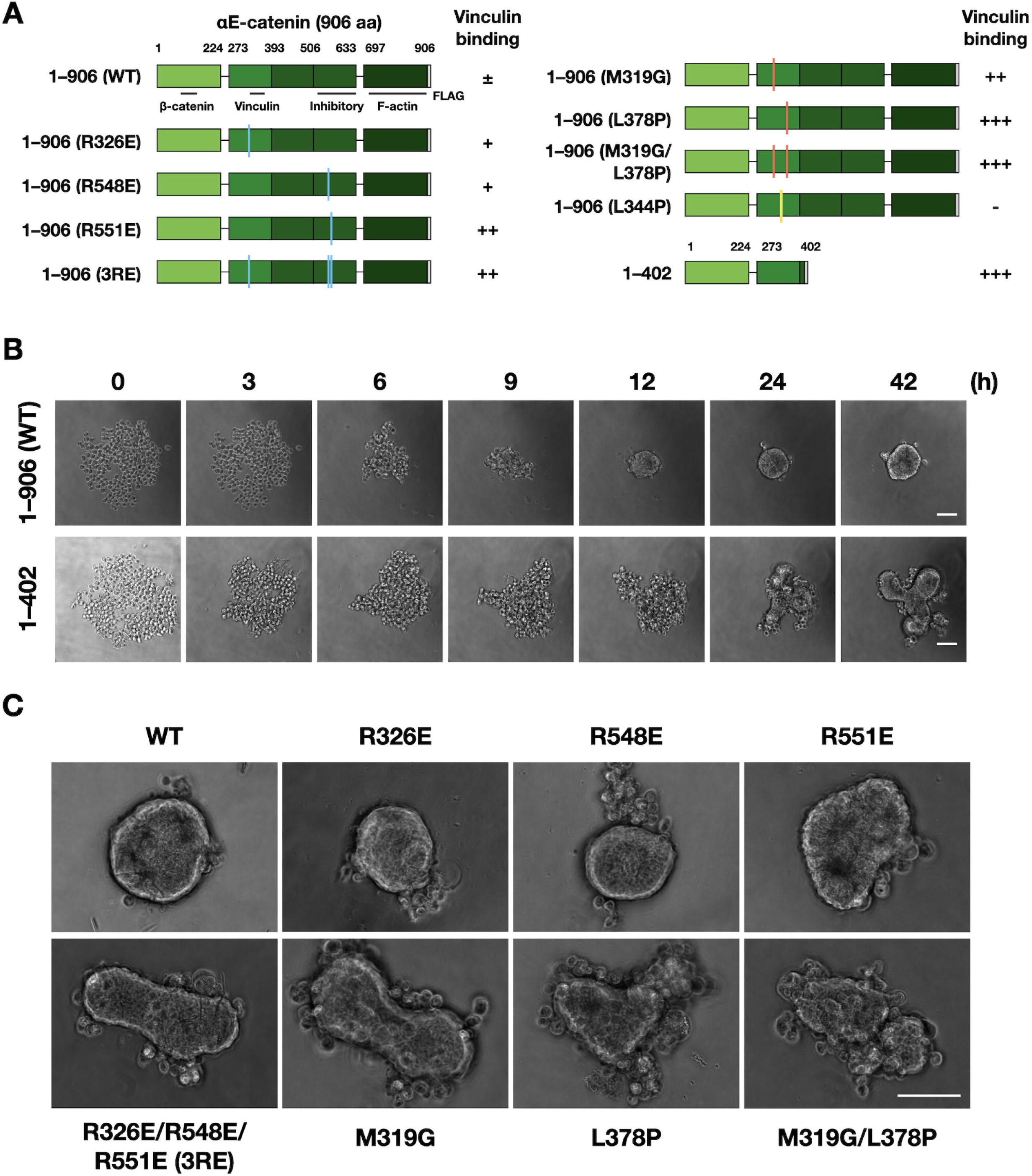
Wild-type α-catenin-expressing cells form round-shaped spheroids in round-bottomed wells, which is not supported by tension sensitivity mutants. **(A)** Schematic drawing of α-catenin and its mutants. The residue numbering for mouse αE-catenin is given. 1–906; full length, 1–402; C-terminus deletion. WT; wild-type, 3RE; R326E/R548E/R551E. Colored lines on the domain illustration indicates the positions of point mutations. Functional domains and strength of vinculin binding affinity are indicated. -; no binding, ±; moderate, +; strong, ++; very strong, +++; extremely strong (Hirano et al., 2018). **(B)** Still images by time-lapse microscopy. R2/7 cells expressing WT (1–906; top) or mutant (1–402; bottom) α-catenin were seeded on round-bottomed wells, respectively, and live-imaged for 42 h. **(C)** Spheroids of various mutants after 48 h culture. Scale bar, 100 µm.

First, spheroid formation experiments were conducted in commercially available nonadherent round-bottomed well plates to confirm if the tension sensitivity of α-catenin is involved in epithelial morphogenesis (**Fig. 1**, **Movie 1, 2**). R2/7 cells stably expressing WT and mutant α-catenin (**Fig. 1A**) were seeded into round-bottomed wells, and the morphology of the cell mass was recorded over time. Cells expressing WT α-catenin adhered to each other to form a single aggregate, which then gradually deformed into a spherical shape with a smooth surface (**Fig. 1B**, **Movie 1**). In contrast, cells expressing the C-terminal deletion mutant (1-402) similarly formed a single aggregate but with a distorted shape (**Fig. 1B**, **Movie 1**). Point mutants with different levels of increased tension sensitivity (R326E, R548E, R551E, R326/548/551E; 3RE, M319G, L378P, and M319G/L378P) (41) expressing cells also formed distorted spheroids according to the degree of enhanced tension sensitivity (**Fig. 1C**, **Movie 2**). Therefore, when α-catenin tension sensitivity is normal, cells form round spheroids in this system, while rounding is suppressed when the tension sensitivity is increased, indicating the importance of α-catenin tension sensitivity in spheroid morphogenesis into a round shape. It is easily assumed that clonal cells with the same genetic background would form symmetrical round spheroids. In this point of view, formation of deformed spheroids by cells expressing α-catenin hypersensitive mutants is unexpected and very intriguing.

### Construction of an experimental epithelial morphogenesis model and quantitative analysis

In the round-bottomed wells, the projection image of the initial shape from above was circular similar to the final shape, which was hard to control (**Fig. 1**, **Fig. 2A**). Therefore, developing a new 3D culture model was attempted in which the initial shape could be controlled, and quantitative image analysis is possible. Dispersed cells were seeded into V-bottom rectangular microwells made of nonadhesive polydimethylsiloxane (PDMS) molded in a metal mold (**Fig. 2A**).

**Fig. 2.**
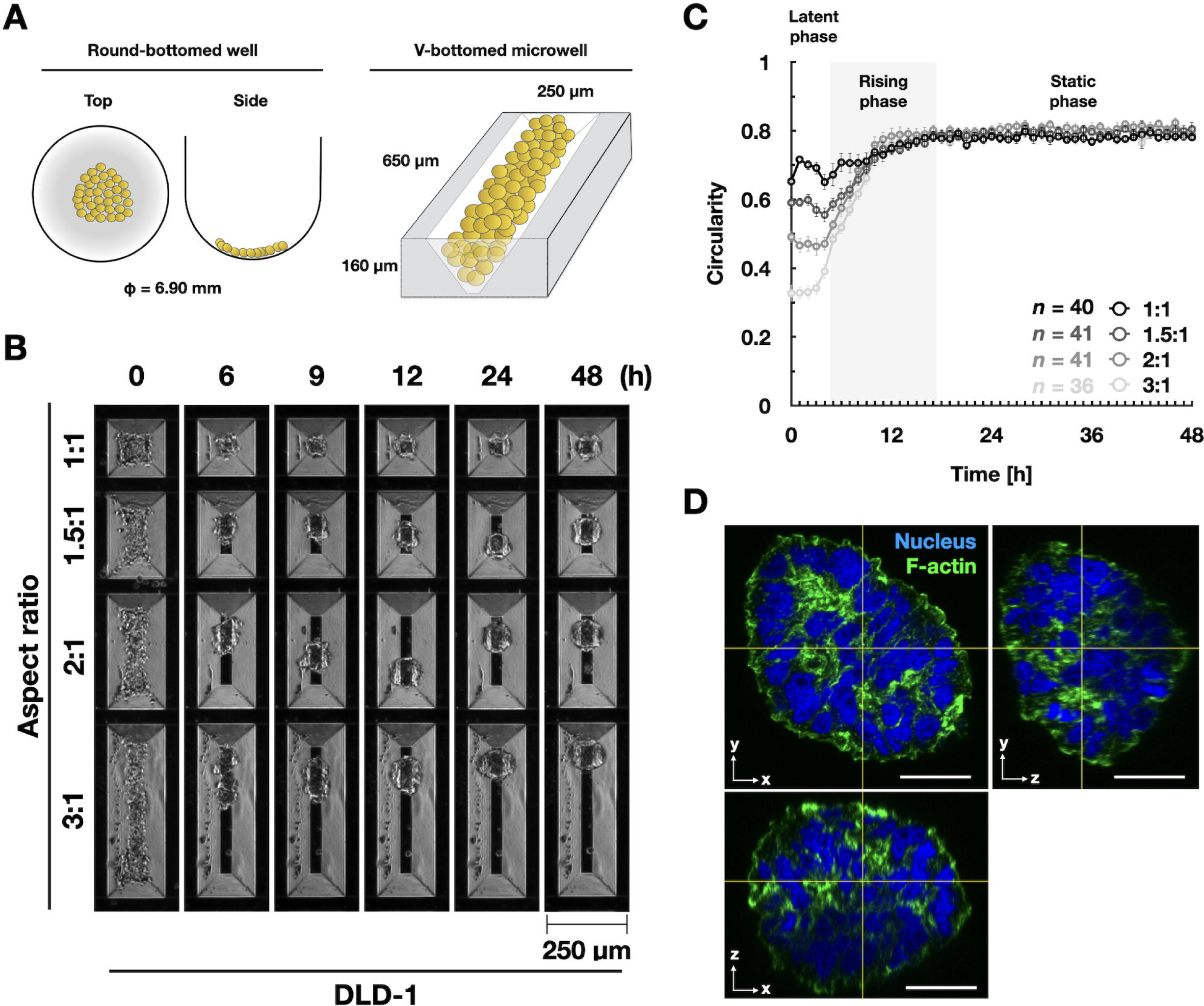
Initial aspect ratios of cell masses and final spheroid circularity. **(A)** Schematic drawings of a round-bottomed well and a V-bottomed microwell. (**B**) Formation of round spheroids in V-bottomed microwells. Still images by time-lapse microscopy. Scale bar, 250 µm. DLD-1 cells were seeded on V-bottomed microwells which have 1:1, 1.5:1, 2:1 or 3:1 aspect ratio, respectively, and live-imaged for 48 h. **(C)** The circularity of spheroids that measured every 1h. Error bars show mean ± 95%CI. (**D**) Visualization of nucleus and F-actin of a round spheroid. R2/7 cells expressing WT α-catenin were seeded on V-bottomed microwells which have 3:1 aspect ratio, cultured for 48 h, fixed, stained, and imaged. The XZ and YZ view images were produced at the horizontal and vertical lines indicated in the XY view image, respectively, showing 3D round morphology. Scale bar, 20 µm.

To investigate the relationship between the initial and final shapes of the cell aggregates, spheroid formation experiments with DLD-1 cells were performed using the microwell with several different aspect ratios (**Fig. 2**, **Movie 3**). The initial shape of the cell mass remained unchanged until about 4–5 h after seeding and changed to a spherical shape by 12–18 h (**Fig. 2B**, **Movie 3**). After that, no change in shape was observed (**Fig. 2B**, **D, Movie 3**). To evaluate this temporal change more objectively, the projected contour of the spheroid in the XY plane was traced and the temporal change of its circularity was plotted (**Fig. 2C**). Although the cells were actively oscillating up to 4–5 h after seeding, there was little change in the circularity (the *latent phase* in this report). An increase in circularity was observed between 5 and 18 h, especially up to 12 h (*rising phase*). After that, the circularity remained almost constant (*static phase*). The final circularity and the length of time to reach it were almost constant regardless of the aspect ratio of the initial shape. Thus, establishing an experimental model that can quantitatively capture the morphological characteristics of cell aggregate (spheroid) formation was successful. In this suspension culture system, it was found that DLD-1 cells eventually became spherical even if the aspect ratio of the initial shape was large up to 3:1.

Next, the dependence of this morphogenetic process on α-catenin was confirmed (**Fig. S1, Movie 4**). As shown in **Fig. 2**, DLD-1 cell aggregates formed round spheroids (**Fig. S1A–C, Movie 4**). In contrast, R2/7 cells lacking α-catenin expression showed no change in cell mass shape over time. Stable expression of full-length WT α-catenin (1-906) rescued this abnormality fully (**Fig. S1A–C, Movie 4**). Although other proteins may be involved in cell adhesion in general, this morphogenesis depends on the presence of the proper cadherin-catenin complex containing α-catenin.

### Detailed analyses of α-catenin tension sensitivity in the process of rounding morphogenesis

First, the effect of mutations that increase the tension sensitivity of α-catenin on the spheroid formation was examined (**Fig. 3**, **Movie 5**). The triple point mutations, 3RE, in which tension sensitivity is increased compared to WT, caused a delay in the shape change, although the final morphology of the spheroid was similar to that of WT (**Fig. 3A**, **Movie 5**). Cells expressing mutant α-catenin with hypersensitivity to tension (L378P or the C-terminal deletion, 1–402) showed insufficient rounding and less smooth contours than WT, respectively. Although the onset of the rising phase was similar (5–6 h after seeding), the slope of the increase in circularity was smaller than that of the WT (**Fig. 3B**), and the circularity remained lower than that of the WT for 48 h (**Fig. 3C**). We visualized cell-cell adhesion in spheroids after 48 h using immunofluorescence staining. TJs were formed in a typical uniform network in WT-expressing cells indicating that the cortical cell layer of the spheroid is considered a fully developed epithelial sheet (**Fig. 3D**). Neither the C-terminal deletion nor point mutations (3RE and L378P) affected the formation of TJs (**Fig. 3D**). These results quantitatively demonstrate that appropriate tension sensitivity of α-catenin is vital for rounding during spheroid formation. The morphogenetic abnormality caused by increased sensitivity is not due to a severe disturbance in the formation of cell-cell adhesion.

**Fig. 3.**
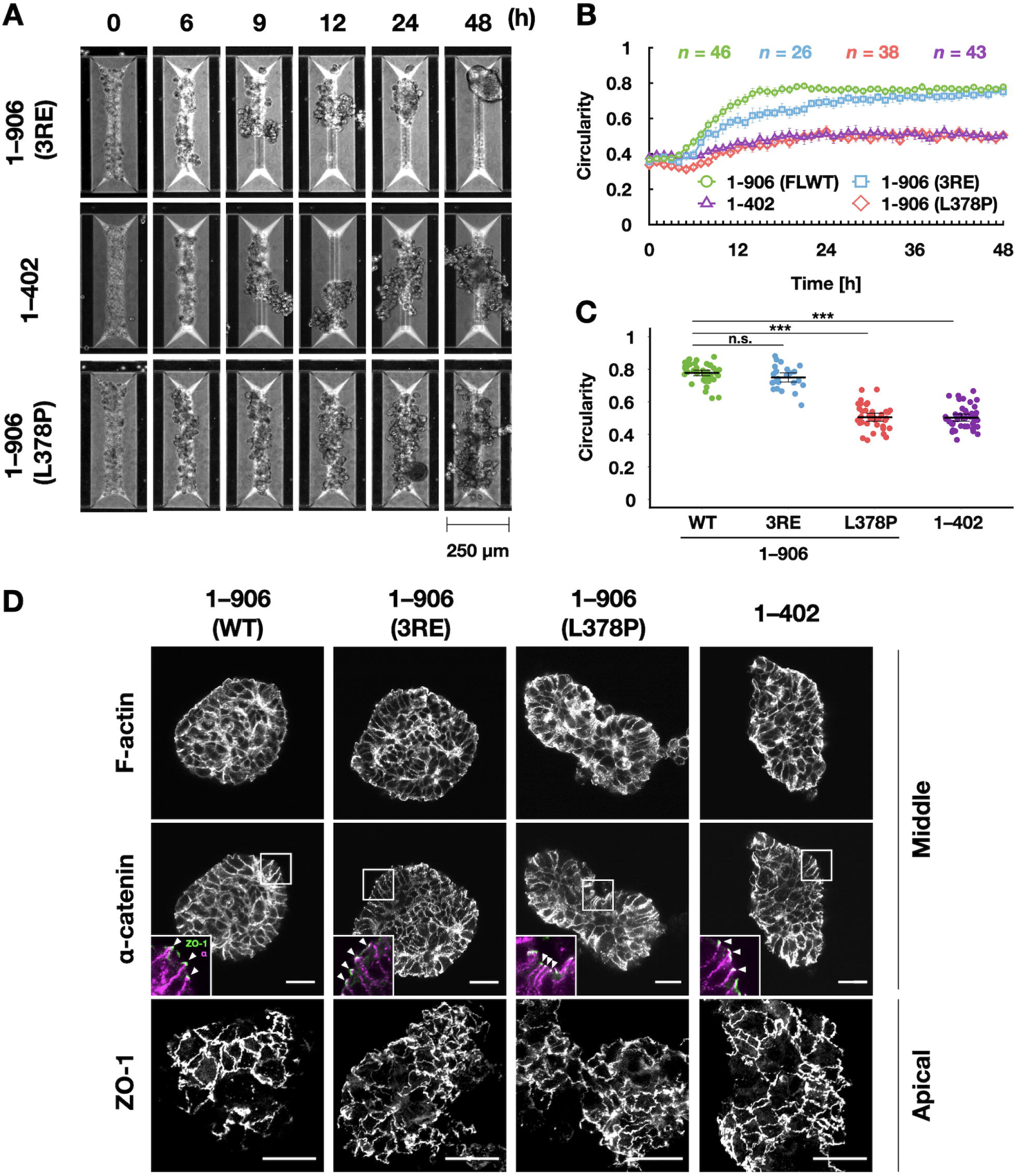
Increased tension sensitivity of α-catenin suppresses increase in circularity of spheroids in V-bottomed microwells. **(A)** Still images by time-lapse microscopy. Scale bar, 250 µm. R2/7 cells expressing α-catenin mutants with increased sensitivity [1–906 (3RE), top, 1–402, middle, 1–906 (L378P), bottom] were seeded on V-bottomed microwells, respectively, and live-imaged for 48 h. **(B)** The circularity of spheroid contour was measured and plotted against time. Error bars show mean ± 95%CI. **(C)** The circularity of spheroids at 48 h after seeding. Hypersensitive mutations suppress increase in circularity. Error bars show mean ± 95%CI. (***; *P*<0.001, n.s.; not significant.) **(D)** Visualization of F-actin, FLAG-tagged α-catenin and ZO-1, showing proper distribution of F-actin and α-catenin together with tight junction networks revealed with ZO-1. White arrowheads in inset indicate apical cell-cell junctions. Cells were cultured on V-bottomed microwells for 48 h, fixed, stained, and imaged. ZO-1 images in the bottom panels are maximum intensity projection of 20 slices (20 µm in depth) from the apical surface of spheroids. Display range of pixel intensity was adjusted independently to clarify junctional structures. ZO-1, green; α-catenin, magenta(inset). Scale bars, 20 µm.

The vinculin binding-null mutant (L344P; **Fig. 4**) (46) can be interpreted as a dull tension sensitivity mutant that cannot respond to any degree of tension for vinculin binding. This mutation caused a slight delay in the increase in circularity but did form round spheroids (**Fig. 4A**, **B, Movie 5**). The circularity during the static phase was significantly higher in cells expressing α-catenin (L344P) than in WT-expressing cells (**Fig. 4C**). Cells expressing this mutation also exhibited normal intercellular adhesion formation (**Fig. 4D**), as did the WT and other mutants (**Fig. 3D**). These results led us to think that rounding morphogenesis in this system requires very low tension, recruiting less vinculin to AJs.

**Fig. 4.**
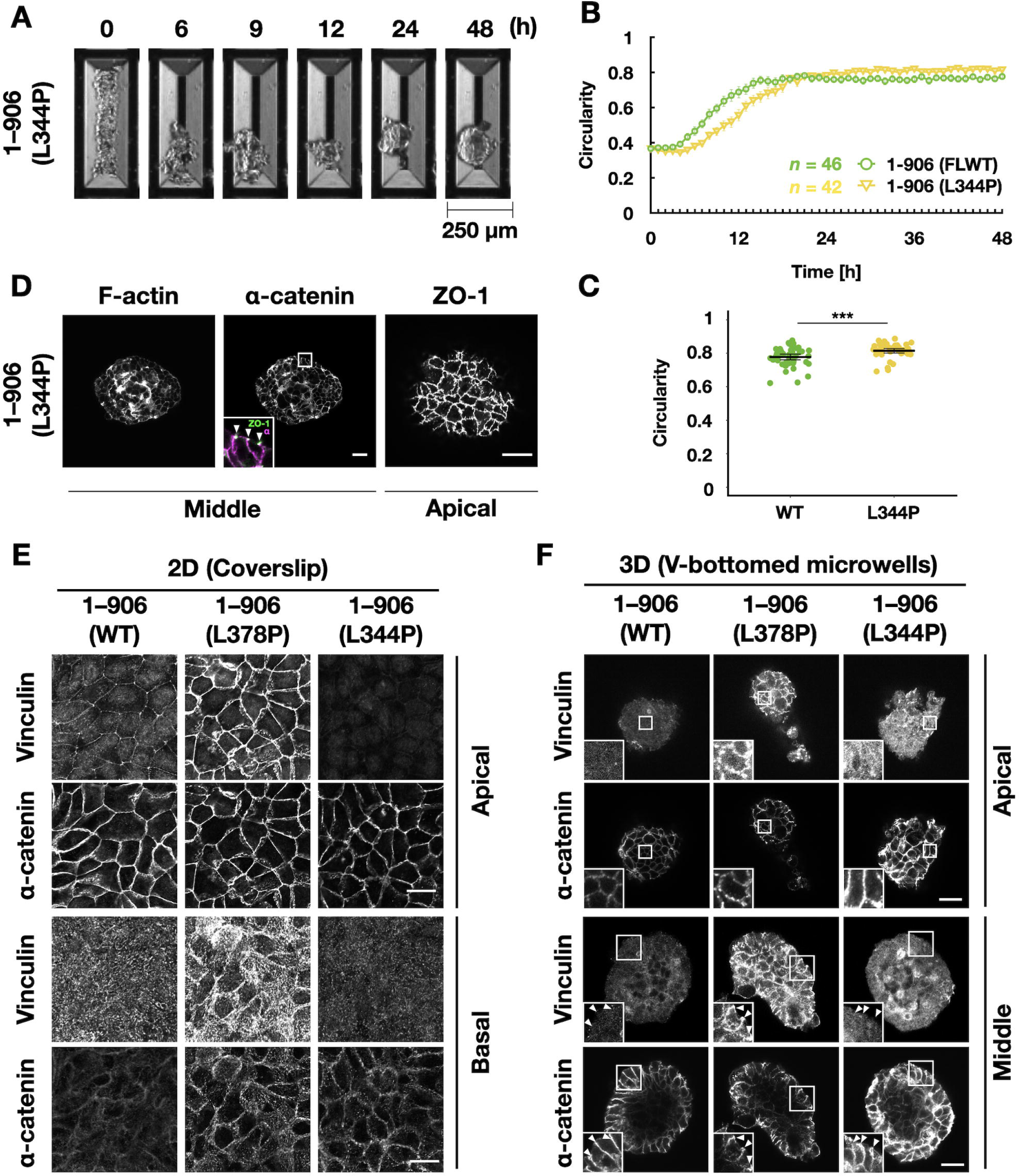
Vinculin does not accumulate at AJs during rounding of WT α-catenin-expressing spheroids on V-bottomed microwells. **(A)** Still images by time-lapse microscopy. Scale bar, 250 µm. R2/7 cells expressing dull mutant α-catenin [1–906 (L344P)] that lacks vinculin binding ability were seeded on V-bottomed microwells and live-imaged for 48 h. **(B)** The circularity of spheroid contour was measured and plotted against time. Error bars show mean ± 95%CI. **(C)** The circularity of spheroids at 48 h after seeding. Error bars show mean ± 95%CI. (***; *P*<0.001.) **(D)** Visualization of F-actin, FLAG-tagged α-catenin and ZO-1 showing proper distribution of F-actin and cadherin together with tight junction networks revealed with ZO-1. White arrowheads in inset indicate apical cell-cell junctions. ZO-1, green; α-catenin, magenta (inset). Cells were cultured on V-bottomed microwells for 48 h, fixed, stained, and imaged. **(E, F)** Immunostaining of Vinculin and FLAG-tagged α-catenin. Cells were cultured on coverslips (**E**) or V-bottomed microwells (**F**) for 48 h, fixed, stained, and imaged. In WT spheroids, tension-dependent vinculin accumulation to AJs (white arrowheads) is not obvious, showing less tension applied at AJs. Scale bars, 20 µm.

In 2D culture, WT-expressing cells showed vinculin accumulation mainly at cell boundaries, especially at vertexes where several cells meet (**Fig. 4E**). In contrast, there was little vinculin accumulation at cell boundaries in 3D culture (**Fig. 4F**). The L378P mutation in α-catenin caused substantial vinculin accumulation at the cell boundary in 2D and 3D cultures, while the L344P mutation did not. Cells cultured in the presence of blebbistatin from seeding until 48 h showed a similar spheroid formation to untreated cells (**Fig. S2, Movie 7**), indicating that the contractile force of myosin II does not contribute much to the spheroid formation in this system. When myosin II activity was suppressed in cells expressing point mutants (3RE and L378P), there was no obvious difference in morphogenesis compared with untreated cells, indicating that myosin II activity is not involved in morphological abnormalities caused by the mutations. (**Fig. S2, Movie 7)**.

Alternatively, when actin polymerization and depolymerization were inhibited by cytochalasin D (CytoD) and Jasplakinolide (Jasp), respectively, both inhibitors completely suppressed round spheroid formation (**Fig. S3, Movie 8**). It clearly indicates that the round spheroid formation is strongly dependent on actin remodeling. Dynamic actin remodeling is needed for cell motility (51). The results can also be interpreted as the association of F-actin with the cadherin-catenin complex is necessary for morphogenesis, which is consistent with previous reports using α-catenin lacking the actin-binding domain in mice (33) and *Drosophila* (28).

### The effect of tension sensitivity of α-catenin on the maturation process of AJ

Since a protein molecule α-catenin is an important constituent of the protein complex, AJ, it was hypothesized that abnormal tension sensitivity might affect the process of AJ formation and suppress rounding. To obtain high-resolution images, cell-cell adhesion formation after seeding on coverslips was examined (**Fig. 5**). As previously observed (52), cells formed punctate AJs (punctum adherens; PAs) at the early stages after seeding (9–12 h), and then they were reorganized into linear adhesions (zonula adherens; ZAs) (**Fig. 5A**). The calcium switch experiment was also performed. Broken intercellular adhesions by removal of calcium from the culture medium were reformed by replenishing calcium (**Fig. S4**). Furthermore, the morphology of PAs stably found at the peripheries of small cell islands was compared (**Fig. S5**). In total, α-catenin (L344P)-expressing cells formed longer and more PAs, while the length and number of PAs were reduced in α-catenin (L378P)-expressing cells (**Fig. 5A, B, C; Fig. S4A, B, C; Fig. S5**). The L378P mutation also caused TJs to form earlier (**Fig. 5D; Fig. S4D**). Since cells seeded on the coverslips first form PAs, which are then transformed into ZA (52), these results suggest that increased tension sensitivity of α-catenin promotes the maturation of cell-cell adhesion or causes ZA formation, skipping the PA formation process.

**Fig. 5.**
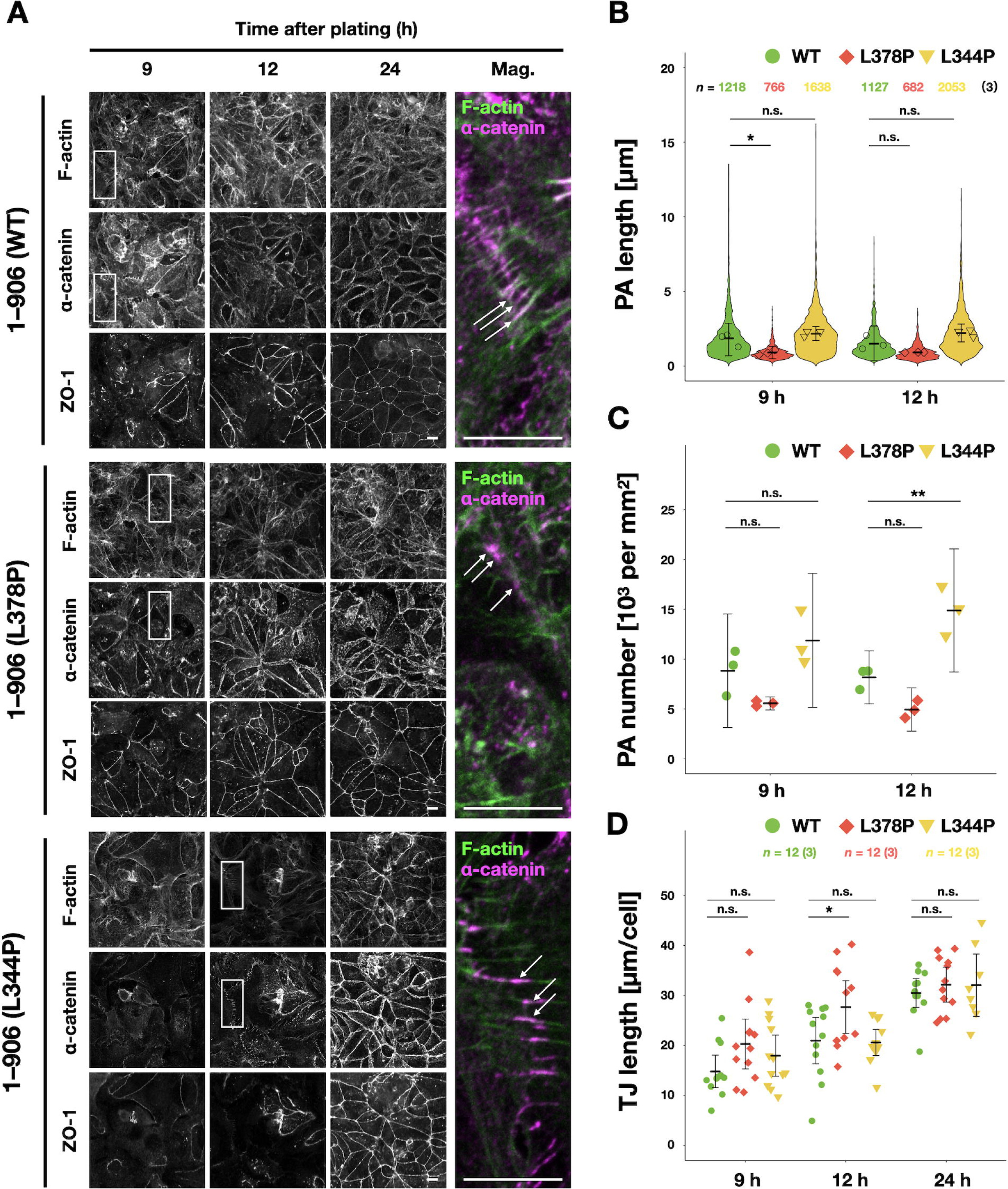
Tension sensitivity mutation of α-catenin alters junctional maturation. **(A)** Visualization of FLAG-tagged α-catenin (magenta) and ZO-1, and of F-actin (green). R2/7 cells expressing wild-type 1–906 (WT; top), hypersensitive mutant 1–906 (L378P; middle), or dull mutant 1–906 (L344P; bottom) α-catenin were seeded on coverslips, respectively, cultured for 9, 12, 24 h, fixed and stained. White arrows indicate PAs. Scale bars, 20 µm. **(B)** Measurement of PA length. Violin plots show data distribution and each dots represents mean values of each biological replicates. **(C)** Measurement of PA number normalized by area. Each dots represents mean values of biological replicates. **(D)** Measurement of TJ length revealed by ZO-1 normalized by cell number over time, showing fast maturation of TJ network in the hypersensitive mutant (L378P). Each dots represents technical replicates. (**B–D**) Error bars show mean ± 95%CI. (*; *P*<0.05, **; *P*<0.01, n.s.; not significant.) See *Materials and Methods* for details of the measurements.

There was an attempt to analyze the process of cell-cell adhesion formation in suspension culture. However, it was tough to prepare fixed samples for observation because cell aggregates in the early stages were easily broken by gentle pipetting. To investigate the formation of cell-cell adhesion in 3D morphogenesis, cell aggregates consisting of a small number of cells on Matrigel were formed (**Fig. S6**). No obvious PA adhesion was observed even at early time points (6–9 h) (**Fig. S6**). Since the formation of PAs is tension-dependent (30) and since it is likely that AJs are not subjected to strong forces during the spheroid formation (**Fig. 4F**, **S2**), it is likely that PAs do not fully develop under such conditions.

### The effect of membrane internalization on epithelial cell morphogenesis

The turnover of component molecules is fast in PAs, or similar spot-like cadherin-based adhesions, and relatively slow in ZAs (53, 54), suggesting that PAs are more plastic than ZAs. Although there was no clear PA formation in 3D culture, it was speculated that AJs at the early stages during maturation were formed during the round spheroid formation. Decreased AJ plasticity causes abnormalities in tissue rearrangements such as cell intercalation (55). Based on these findings, it was hypothesized that AJ plasticity is intimately involved in the round spheroid formation. AJ plasticity is known to be regulated by the internalization of membrane proteins, including cadherins, and dynamin GTPase mediates E-cadherin endocytosis (56, 57). The effect of dynamin inhibition by dynasore on rounding of spheroids was examined (**Fig. 6**, **Movie 9**). Dynamin inhibition caused suppression of rounding for both WT α-catenin-expressing and mutant-expressing cells (**Fig. 6A**–**C, Movie 9**). These results support the hypothesis.

**Fig. 6.**
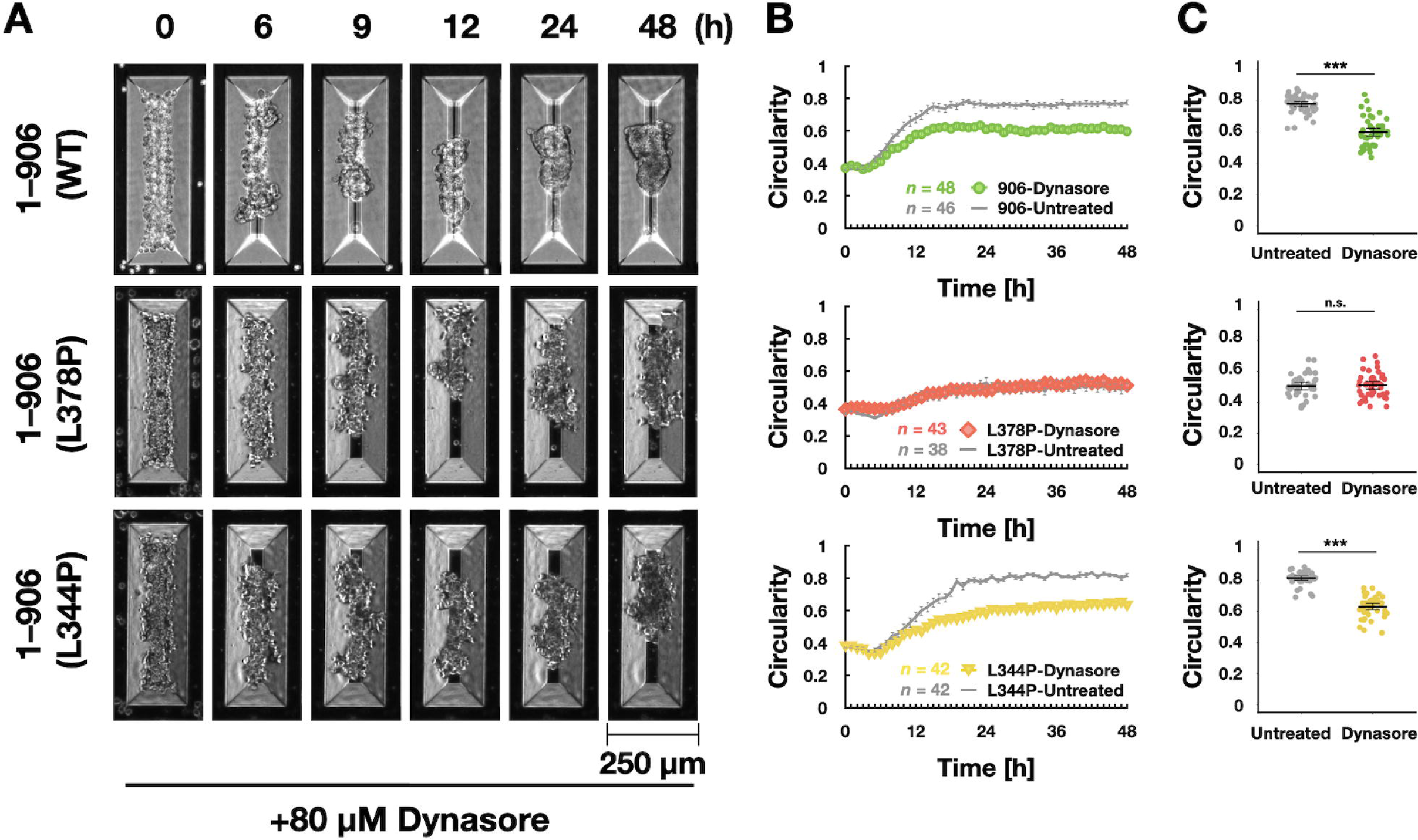
Dynamin inhibition suppresses increase in circularity of spheroids formed in V-bottomed microwells. **(A)** Still images by time-lapse microscopy. Scale bar, 250 µm. R2/7 cells expressing wild-type (1–906 (WT; top), hypersensitive mutant [1–906 (L378P; middle)], or dull mutant [1–906 (L344P; bottom)] α-catenin were seeded on V-bottomed microwells, respectively, and live-imaged for 48 h in the presence of 80 µM Dynasore, a dynamin inhibitor. **(B)** The circularity of spheroid contour was measured and plotted against time. Error bars show mean ± 95%CI. **(C)** The circularity of spheroids at 48 h after seeding. Note that in L378P mutant, decrease in circularity by dynasore treatment was not detected. Error bars show mean ± 95%CI. (***; *P*<0.001, n.s.; not significant.)

### Increased tension sensitivity of α-catenin inhibits collective rather than local cellular rearrangements during round spheroid formation

The behavior of individual cells during rounding was then analyzed based on the idea that rounding should not occur unless the spatial arrangement of cells is changed. For this cell rearrangement in an epithelial sheet to occur, AJ remodeling with dissociation of old AJs and formation of new AJs is necessary. First, to test the hypothesis that the displacement or rearrangement of neighboring cells leads to round spheroid formation, cell nuclei using GFP-tagged Histone 2B were visualized and cell movements during the spheroid formation was analyzed (**Fig. 7, Movie 1 0,11)**. To obtain better Z-stack images by confocal microscopy, we utilized amorphous fluoropolymer CYTOP, which has low refractive index near to water, as a substrate of the microwells. Here the focus was on the rising phase (6–18 h after seeding) and nuclei positions were tracked (**Fig. 7A–E**). By calculating the correlation coefficients (*r*) of the direction of displacement of each cell of a pair of neighboring cells in each frame, we can determine whether the two cells are moving in the same direction with a high positive correlation (0 < *r* ≦ +1) or away from each other with high negative correlation (0 > *r* ≧ -1) (**Fig. 7A**, **Movie 1 0**). The distribution of *r* over the entire period shows two peaks, one near 0 and the other near +1, indicating that the neighboring cells often move in a highly correlated or uncorrelated manner (**Fig. 7B**). Events, which cells pass by each other with negative *r* do not occur very often. Next, the changes in the distribution of *r* over time in a 2D density plot with time as the horizontal axis was visualized (**Fig. 7C**). In the two conditions in which rounding proceeded normally (WT and L344P mutant α-catenin-expressing cells), the proportion of highly correlated movements tended to increase during the first half of the rising phase (6–12 h, especially up to 9 h), when the change in circularity was most remarkable (**Fig. 7C**). The distribution of r was almost the same in L378P mutant α-catenin-expressing cells that cannot form round spheroids (**Fig. 7B**). Furthermore, the inhibition of dynamin had almost no effect on these cell movements (**Fig. 7B**). Under these conditions, there was little change in the distribution over time (**Fig. 7C**). There was no difference in the percentage of highly correlated movements even when the range was limited to 6–9 h (**Fig. 7D**, **E**). These results indicate that this rounding process cannot be explained mainly by local exchanges of neighboring cell positions, detected as movements with negative *r*. Thus, the hypothesis is denied.

**Fig. 7.**
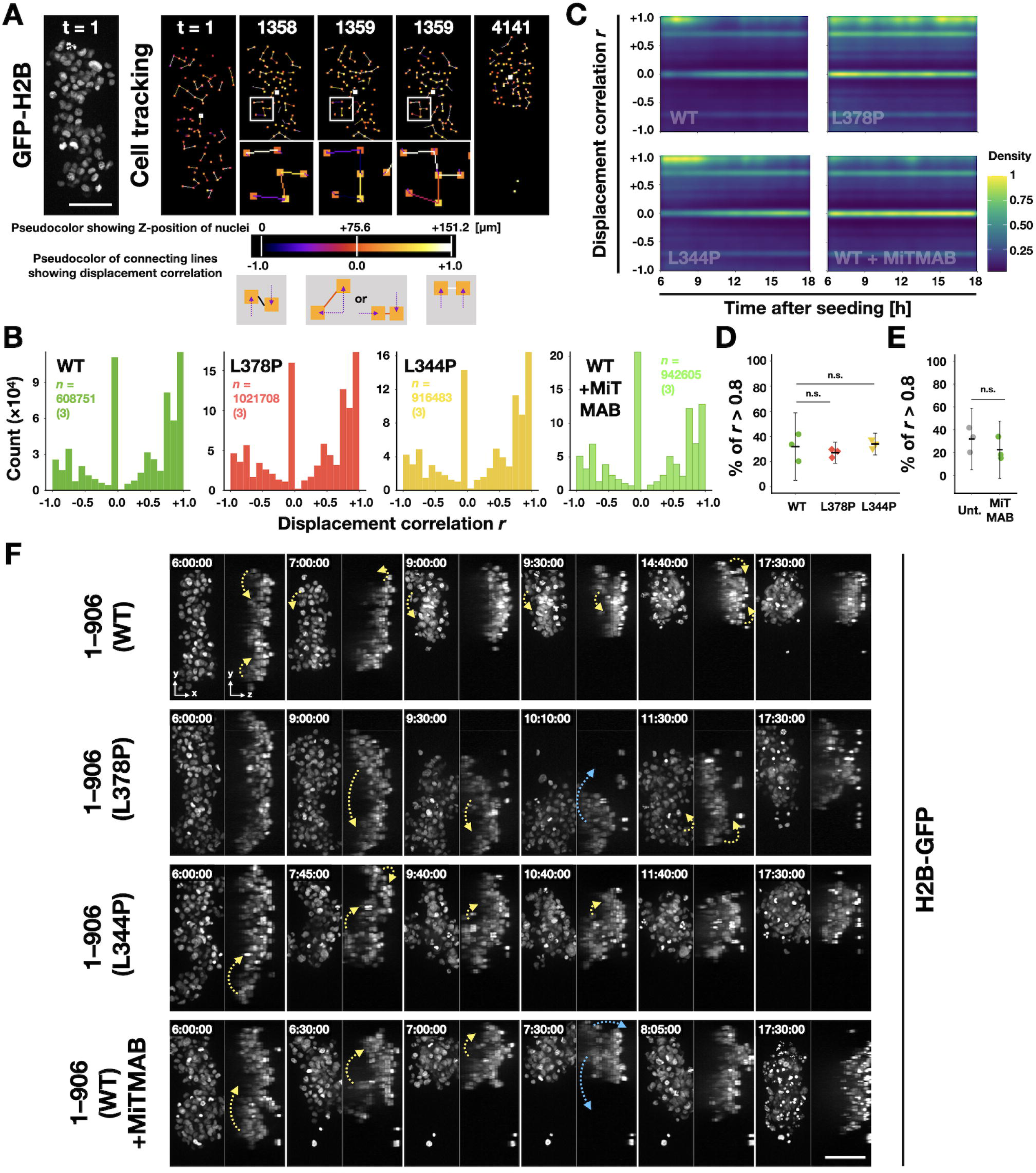
Collective but not single cell rearrangement is found often in the spheroid rounding. **(A)** A GFP-H2B image showing nuclei (left) representing the position of cells and visualization of cell tracking analysis process (right). Each cell position in the XY plane is shown in small square and its color expresses the depth (‘colder’ color shows deeper (closer to bottom) position). Displacement of each cell was calculated by comparing its position to that of the previous frame. Then, displacement correlation (*r*) was calculated by comparing 3D displacement vectors of two neighboring cells (connected by lines) and shown by the same color scale (‘colder’ color shows negative, ‘hotter’ color shows a positive correlation, respectively). Frame numbers t are indicated in each image (frame interval = 10 sec). R2/7 cells expressing H2B-GFP and wild-type [1–906 (WT)] or mutants [1–906 (L378P), or 1–906 (L344P)] α-catenin were seeded on V-bottomed microwells, respectively, cultured for 6 h and then live-imaged for 11.5 h. R2/7 cells expressing H2B-GFP and wild-type α-catenin were also seeded and cultured as well and then live-imaged for the same duration in the presence of 10 µM MiTMAB, a dynamin inhibitor (WT+MiTMAB). **(B)** Histogram showing data distribution of displacement correlation *r*. **(C)** Two-dimensional density plot showing data distribution of displacement correlation *r* over time. The density is normalized to 0–1 and coded by pseudo color. **(D, E)** Percentages of cell displacement pairs *r* greater than 0.8, at between 6–9 hours after seeding. Each dot represents a biological replicate. Error bars show mean ± 95%CI. (n.s.; not significant.) **(F)** Still images by confocal time-lapse microscopy. Stack images are shown as maximum intensity projection images of the top (XY) or transverse (YZ) view. Time after seeding is indicated in each frame. Yellow and blue arrows indicate the direction of cell mass movement (‘folding’ and ‘opening’, respectively). Scale bars, 100 µm.

To search for other features to focus on, we attempted to simultaneously observe cell movement from the overturned plane using the stacked image data obtained in previous experiments (**Fig. 7F**, **S7, Movie 1 1**). As a result, it was observed that in WT α-catenin-expressing cells, long cell aggregates at the initial state repeatedly bent from both ends or at the center and gradually became round (**Fig. 7F** **-WT, S7A, Movie 1 1**). Those cell aggregates appear to constitute several parts. One part comprises few cells adhering to each other. Those parts move as a group of cells like swing arms using the region between two parts as a supporting point. When the parts come into contact with each other by bending, they fuse to form a single component. This event is seen as a jump in circularity during the rising phase (Fig. S7). Whenever this fusion happens, AJ remodeling is required; some of the existing AJs should be broken, and new ones should be formed (**Fig. S8**). The same behavior was observed in cells expressing the L344P mutant (**Fig. 7F** **-L344P, Movie 1 1**). Similar bending motions were observed in cells expressing the L378P mutation and WT cells treated to suppress membrane uptake (**Fig. 7F** **-L378P, and WT + MiTMAB, Movie 1 1**). We note that we used MiTMAB instead of dynasore to inhibit dynamin, because dynasore was turned out to show phototoxic effects during fluorescent imaging. However, they could not hold the bent state and returned to their original elongated shape after a while (**Fig. 7F** **-L378P, and WT + MiTMAB, Movie 1 1**). These results made us imagine that AJ’s low plasticity would suppress such fusion between parts.

To test this, an experiment was performed in which preformed spheroids were fused (**Fig. 8**, **S8, Movie 1 2**). To use spheroids with similar size, relatively small number (about 10–20) of cells were seeded into V-bottomed microwells with an aspect ratio of 1:1, and two spheroids were picked in the static phase (formed after 24 h of culture), and transferred to other V-bottomed microwell plates to achieve contact between them. In WT and L344P mutant α-catenin-expressing cells, the adhesive surface expanded within few hours after contact, forming a single round spheroid (**Fig. 8A**, **Movie 1 2**). In contrast, cells expressing the L378P mutation did not develop a clear contact surface even after 9 h (**Fig. 8A**, **Movie 1 2**). The L378P mutation suppressed the expansion of the contact area and kept the distance between two spheroids long, whereas the WT and L344P mutants expanded the contact surface and shortened the distance between the two spheroids (**Fig. 8B**, **C**). These results show that the appropriate tension sensitivity of α-catenin contributes to the fusion between spheroids, leading to spheroid rounding probably through the AJ remodeling (**Fig. 9**).

**Fig. 8.**
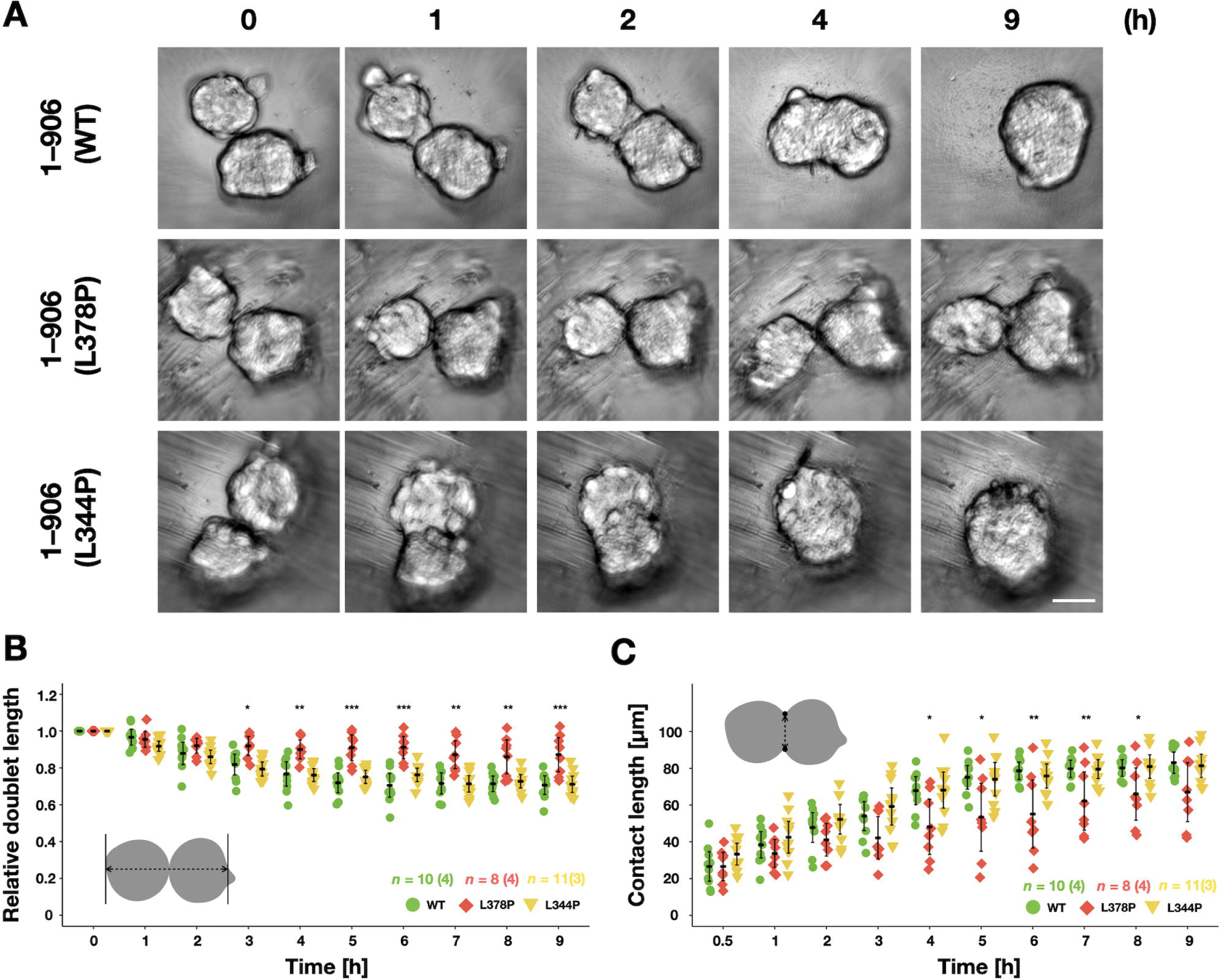
Fusion experiments of spheroids of cells expressing α-catenin with different tension sensitivity. **(A)** Still images by confocal time-lapse microscopy. R2/7 cells expressing wild-type (1–906 (WT; top), hypersensitive mutant (1–906 (L378P; middle)), or dull mutant (1–906 (L344P; bottom)) α-catenin were seeded on V-bottomed microwells, respectively, cultured for 24 h. Spheroids were then transferred onto V-bottomed wells of 96-well plate and live-imaged for 9 h. Hypersensitive mutation (L378P) suppressed spheroid fusion, suggesting the importance of plasticity of AJ in spheroid fusion. Scale bar, 50 µm. **(B)** The length of a spheroid pair along the long axis was measured. Any protrusions on spheroids were ignored, and values were normalized to that at 0 h. **(C)** The length of new contacts between two spheroids was measured as illustrated. (**B, C**) Error bars show mean ± 95%CI. (*; *P*<0.05, **; *P*<0.01, ***; *P*<0.001, n.s.; not significant.)

**Fig. 9.**
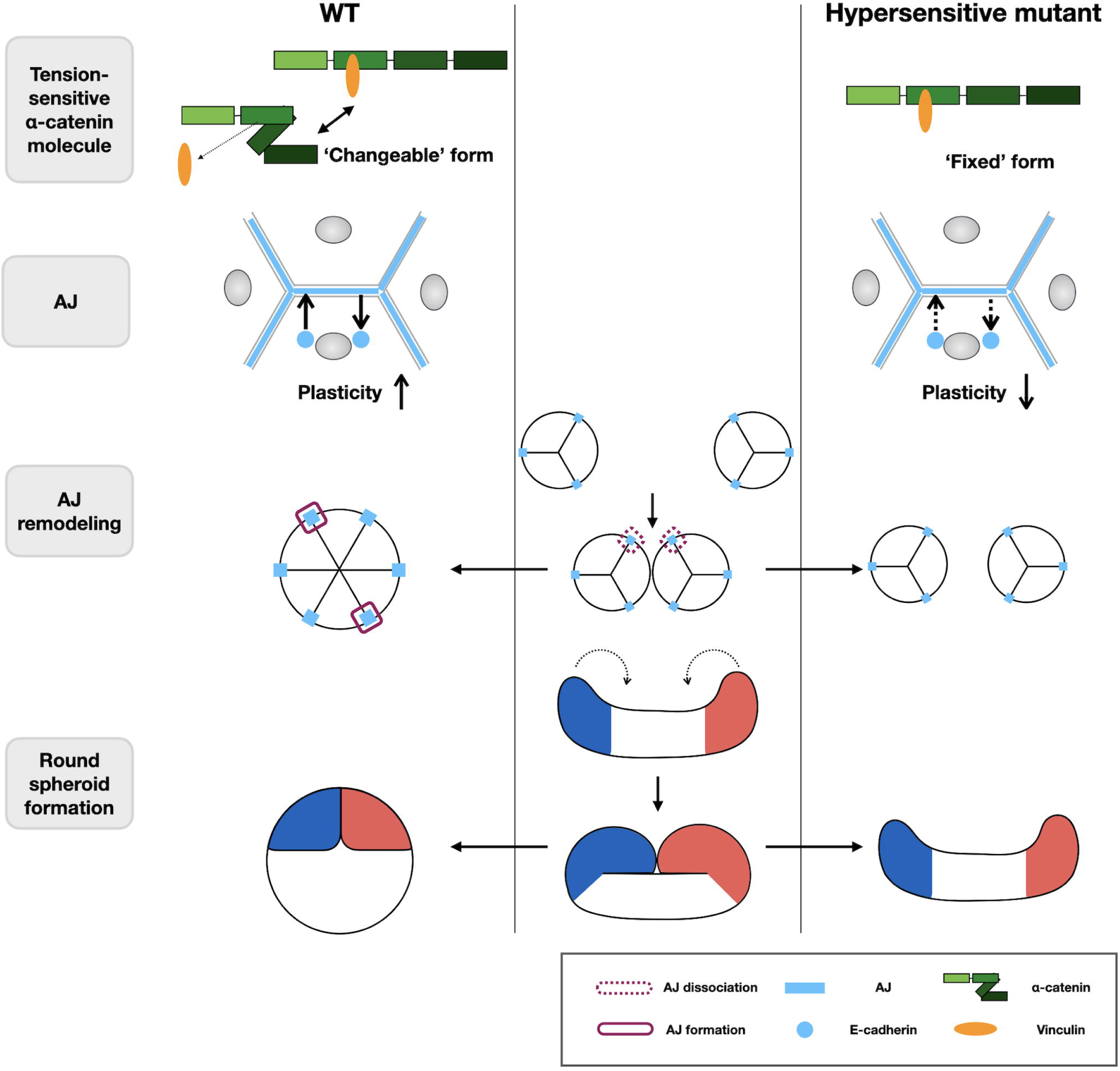
Schematic illustration of the proposed model. Trans-scale relationships from molecular responses to multicellular behaviors during rounding morphogenesis of epithelial spheroid are illustrated. Under suspension condition, elongated cell aggregates gradually deform into spherical shape by folding events. It accompanies AJ remodeling (i.e., dissociation and formation), which requires plasticity of AJs. When α-catenin tension sensitivity is increased to the maximum by mutation, the molecule is fixed to ‘opened’ form, which strongly binds to vinculin. It leads to decrease in AJ plasticity, prevents AJ remodeling, and results in failure of the collective rearrangement. Thus, proper tension sensitivity of α-catenin ensures rounding morphogenesis of epithelial spheroids.

## Discussion

It has been often observed that various tissue culture cells form round spheroids *in vitro*, (6, 7), and many species of organisms exhibit round embryos at early stages (58). In this study, it was found that even rectangular cell masses with high aspect ratios consisting of dissociated cells can transform into round spheroids accompanied by intercellular junction formation. Alternatively, rounding can be regarded as a fundamental property of epithelial cell morphogenesis in suspension culture. The minimum requirements for rounding are as follows: (1) the cells adhere to each other with the same strength, (2) they can change their relative spatial positions. These requirements involve that the adhesion has plasticity with dynamic association and dissociation and that the cells must have motility. It is easy to imagine that a cell mass consisting of individual cells with the same adhesion properties forms a round spheroid when they move randomly to achieve an energetically stable state (7, 59). In contrast, the tissue’s surface tension caused by the cortical tension of individual cells has also attracted attention as an essential factor in driving spheroid’s rounding (7, 59).

There are three phases of spheroid formation (**Figs. 2–4**). There is little change in circularity (**Fig. 2**), but individual cells exhibit oscillatory motions (**Movies 3–6**) at the latent phase. The cell-cell junctions are yet unobserved. Cells after dissociation wait for a sufficient amount of cadherin expression on their surface with repeated weak association and dissociation. In the next rising phase, the circularity increases. TJs are partially observed in this phase, and the cell aggregate compaction begins, suggesting that the AJ formation started at the beginning of this phase. In the third static phase, the circularity does not change, and the TJ network formation is completed on the spheroid surface, suggesting that the strength of cell-cell adhesion exceeds that of cell motility, and the overall shape remains stable (**Figs. 3 and 4**). In this phase, the spheroids can still be seen moving (**Movies 3–6**). Many studies have been conducted to characterize the spheroid formation process over time using factors describing morphology such as diameter and circularity (60-63). However, to the best of our knowledge, this is the first study to define distinct formation stages and analyze the rounding process at the molecular level.

In the rising phase, AJs were thought to be converting from PAs to ZAs, but it was challenging to recognize AJs using immunostaining of spheroids because only a weak force was applied to AJs in spheroids, recruiting a few AJ-specific vinculin. Since the molecular turnover of PAs is faster than that of ZAs (53, 54), the plasticity of AJs in the PA phase is probably high. Since the TJ network adjacent to ZA is also complete by the end of rounding, it is reasonable to assume that AJ remodeling associated with cell rearrangement during rounding occurs primarily during the PA phase. At this stage, however, cell rearrangement by positional exchanges of adjacent cells was not often seen. Cells instead moved collectively as in the gradual folding of a long cell mass (**Fig. 7** **and Movie 1 1**). Even during the static phase, AJ remodeling can occur such that established spheroids could deform and fuse (**Fig. 8**).

In α-catenin mutants with artificially increased tension sensitivity, rounding of spheroids in round-bottomed wells was inhibited with increasing sensitivity (**Fig. 1**). In V-bottomed microwells, the increase in circularity was delayed with moderate sensitivity but eventually reached the same level of circularity as the WT. In the hypersensitive mutants, the increase in circularity was markedly suppressed (**Fig. 3**). To test the possibility that myosin II-induced structural changes in α-catenin and consequent vinculin binding is important for proper rounding, we inhibited myosin II activity with blebbistatin (**Fig. S2**) or vinculin binding using the L344P mutation (**Fig. 4**). In both cases, rounding occurred almost normally. These results show that strong myosin II activity is not required for this morphogenesis. And tension sensitivity for dissociating vinculin under low tension is crucial for rounding morphogenesis in this system. Morphogenesis in this system showed no vinculin accumulation on AJs, even in cells expressing WT α-catenin, suggesting that AJs are not under high tension (**Fig. 4**). Since actin polymerization or depolymerization inhibition does not induce rounding of spheroids at all (**Fig. S3**), cell motility induced by actin polymerization is likely necessary for rounding morphogenesis.

The hypersensitive mutation affects the AJ formation process (**Figs. 5, S4, and S5**), while it supports cell adhesion function enough to form TJ networks (**Fig. 3**). AJs mature into ZAs through the PA stage in the normal process of AJ formation, and TJs are formed along the ZAs. In the hypersensitive mutant, TJs formed without distinct development of PAs and seemed to mature quickly into ZAs. The turnover of AJ components is faster in PA (53, 54) and slower in AJs formed by hypersensitive mutant α-catenin (47). Therefore, it was hypothesized that the tension-sensitive mutation might suppress rounding by reducing AJ plasticity. When general membrane endocytosis proteins, including cadherin, were inhibited using dynasore, a dynamin inhibitor, rounding was also suppressed (**Fig. 6**), supporting the idea that normal turnover of AJ components, or AJ plasticity, is important for rounding morphogenesis.

In addition to cell-cell adhesion, the ability of each cell to change its position is thought to be essential in rounding. The nucleus was tracked for positioning each cell and the direction of displacement of the position of each pair of neighboring cells for each frame was measured. If individual cells were moving randomly, it is expected that the displacement of a significant number of neighboring cells would be indicated as a low correlation toward displacement. But in fact, it was seen only infrequently, with many cells showing displacement in the same direction, i.e., highly correlated movement (**Fig. 7A–E**). Detailed analysis revealed that when some parts of long cell mass bend, a circularity jumps were observed, which was an essential contribution to the increase in circularity (**Figs. 7F** **and S7A, B**). Previous graphs of the increase in circularity (**Figs. 2–4**) showed the average circularity of many spheroids and thus would not have reflected the characteristic morphological changes of individual spheroids. This movement was commonly observed in WT and hypersensitive mutants, but in the WT, the movement was followed by new adhesion and an increase in circularity. In contrast, in the hypersensitive mutant, the bending was not sustained, and the cells moved again, which did not improve circularity. When two cell masses fuse by contacting each other, some of the existing AJs should be broken, and new AJs formed, i.e., AJ remodeling is required (**Fig. S8**). In the WT and mutants with low tension sensitivity, this AJ remodeling appears to occur without problems but in hypersensitive mutants do not. To test this idea, the two spheroids were placed next to each other to see if they would fuse. As expected, only the hypersensitive mutation suppressed spheroid fusion (**Fig. 8**).

α-Catenin has been designed to bind to vinculin when the force on the molecule exceeds a physiologically sufficient magnitude (>5 pN, corresponding to the force generated by a myosin II molecule)(43, 64). Under physiological forces, accumulated vinculin, an actin-binding protein, functions as actin-binding scaffolds in AJs and strengthens them to withstand strong forces (29-31). Suppose this sensitivity set points were higher, i.e., if α-catenin would bind to vinculin under lower tension, smooth remodeling of AJs would not be possible. Thus, evolutionarily, the tension sensitivity of the α-catenin molecule has been optimally tuned.

A typical example of cellular rearrangements has been reported, such as germ-band extension in the *Drosophila* embryo, where local neighbor exchange of cells is vital for morphogenesis (55). In contrast, in this study, such neighbor exchange does not play at least a major role in the rounding of spheroids. *In vivo* epithelial sheets are located on top of the basement membrane, making it very easy for them to move on the substrate. However, spheroids made from epithelial cells alone under experimental suspension culture conditions do not form a basement membrane as seen *in vivo*. The role of cell movement in morphogenesis is thought to be different in cases where cell movement on the basement membrane is easy and in cases where there is no substrate to support cell movement. In the experimental system used in this study, cell motility itself was necessary. In addition, the movement of folding as a group appeared to make a relatively large contribution. The degree of neighbor exchange of cells was low, and their contribution to rounding was probably minor.

α-Catenin reduces AJ plasticity to ensure the force transmission under strong tension and increases plasticity to allow remodeling when the tension is weakened. It is thought that tension sensitivity is essential for successful switching between these two functions. Furthermore, tension range for the switching is also important. In this sense, the α-catenin molecule is considered to be cleverly designed to respond to physiological range of tension. Since the direct actin-binding of α-catenin itself is also tension-sensitive, it is necessary to further investigate the relationship between this and the actual adhesion to substrates *in vivo*.

When cell clones with the same properties gather, it is very natural to think that they form a round aggregate. It is also assumed that many distorted aggregates can be formed by stochastic fluctuations of each cell behavior. In reality, clonal epithelial cells form round spheroids with high probability *in vitro,* and round embryos are formed from eggs with high precision during development. Tension sensitivity of α-catenin appears to support designed morphogenesis protecting from stochastic fluctuations.

### Experimental procedures

#### Cell culture

DLD-1 cells from ATCC, its subclone R2/7 cells (36) (provided by F. van Roy, Ghent University, Belgium), and R2/7 cell lines expressing various α-catenin mutants were cultured in Dulbecco’s Modified Eagle Medium (DMEM; Wako) supplemented with 10% fetal bovine serum (FBS).

### Spheroid formation

DLD-1 cells, R2/7 cells, or R2/7 cell lines expressing WT or mutant α-catenin were cultured on the following containers. For the spheroid formation in round-bottomed wells, nonadherent round-bottomed 96-well plates (EZ-BindShut SP, Iwaki) were used. Cells were suspended in a culture medium, seeded in each well at 100 cells/well, and cultured for 48 h. For the spheroid formation in V-bottomed rectangular microwells, polydimethylsiloxane (PDMS; Sylgard 184, Dow Corning) or CYTOP (CTX-809SP2, Asahi Glass) was shaped by microfabricated metal mold (described in the following subsection) according to manufacture instructions and attached to a 35mm glass-bottomed dish (Matsunami Glass), then treated with 10% Poloxamer 188 (Pluronic F68, GE Healthcare) overnight to prevent cell attachment (65, 66). The dish was rinsed three times using PBS and filled with culture medium, or imaging medium (FluoBrite DMEM, Thermo Fisher) supplemented with 10% FBS. A glass cloning cylinder (inner diameter 7 mm; Iwaki) was placed on the substrate to limit the area of cell seeding, and cell suspension (1 × 10^4^ cells for the cell tracking and 2 × 10^4^ cells for all other experiments) was added to the substrate using. Cells were cultured in the microwells for 48 h avoiding any stream of culture medium that may bring cells out of the wells. For spheroid formation on an ECM gel, 10 µL Matrigel (Corning) was placed on round coverslips (diameter 14 mm, Matsunami Glass) and gelled by incubating for 10 min at 37°C. Cell suspension (1 × 10^5^ cells) was added to the gel, and cells were cultured for 9–24 h.

### Spheroid fusion

For spheroid fusion assays, R2/7 cell lines expressing WT or mutant α-catenin were cultured on V-bottomed square microwells (width 250 µm × height 250 µm × depth 160 µm) for 24 h to obtain relatively small spheroids. Two spheroids were picked up using a micropipette and transferred together into one well of nonadherent V-bottomed 96-well plates (EZ-BindShut SP, Iwaki). Then, the two spheroids were allowed to contact with each other and were cultured for 9 h.

### Preparation of metal mold

The mold for making PDMS microwells was manufactured by ultraprecision cutting using a single crystal diamond tool with an ultrahigh precision machine (NPIC-M200, Nagase Integrex Co., Ltd.). The material of the mold was nickel-phosphate electroless plating on a stainless steel base with diameter of 10 mm and height of 6 mm. The thickness of nickel-phosphate plating was 300 µm. The diamond cutting tool had rectangular edge with flat end of 50 µm and nose angle of 53 degree to form the shape of well wall and bottom. The final depth of cut was 0.4 µm and surface roughness of 4 nm in average was obtained. After the mold was formed by cutting, a fluorocarbon coating was deposited by reactive ion etching machine so that PDMS can be easily removed from the mold.

### Antibodies and reagents

The following primary antibodies were used: anti-α-catenin rabbit polyclonal antibody (C2018) and anti-vinculin mouse monoclonal antibody (VIN-11-5; Sigma-Aldrich); anti-DDDDK tag rabbit polyclonal antibody, which recognizes the Flag tag (MBL); anti-ZO-1 mouse monoclonal antibody (T8-754; a gift from Sa. Tsukita, Osaka University, Japan). Alexa Fluor 488-conjugated donkey anti-mouse IgG antibody, Alexa Fluor 555-conjugated donkey anti-rabbit IgG, or Alexa Fluor 647-conjugated donkey anti-mouse IgG antibodies (Thermo Fisher) were used as secondary antibodies for immunofluorescent staining. Alexa Fluor 488-conjugated or Alexa Fluor 647-conjugated phalloidin (Thermo Fisher) was used for staining actin filaments. 4′,6-diamidino-2-phenylindol (DAPI; Dojindo Laboratories) was used for staining of nuclei.

(−)-Blebbistatin (Wako) was prepared as a 50 mM stock in dimethylsulfoxide (DMSO) and used at 50 µM. Cytochalasin D (Sigma) was prepared as a 2 mM stock in DMSO and used at 2 µM. Jasplakinolide (Calbiochem) was prepared as a 1 mM stock in DMSO and used at 10 µM. Dynasore (Santa Cruz) was prepared as an 80 mM stock in DMSO and used at 80 µM. MiTMAB (Abcam) was prepared as a 100 mM stock in water and used at 30 µM.

### Immunostaining

For immunofluorescence staining, spheroids formed in V-bottomed microwells were collected in microtubes, fixed with 1% paraformaldehyde (PFA) in 100 mM HEPES-KOH (pH 7.5) at room temperature for 30 min, washed once with 30 mM Glycine in PBS (G-PBS) for 10 min, and permeabilized using 0.5% Triton X-100 in G-PBS for 10 min. They were washed once using G-PBS for 10 min and incubated with 4% normal donkey serum (NDS) in G-PBS for 30 min to reduce nonspecific binding of antibodies. They were incubated with primary antibodies overnight at 4°C, washed thrice with G-PBS for 10 min, incubated with 4% NDS in G-PBS for 10 min, secondary antibodies, and fluorescent reagents (fluorescent labeled-phalloidin and DAPI) for 2 h, and washed thrice using G-PBS for 10 min. They were then suspended in Prolong Glass antifade mountant (Thermo Fisher) and cured for 2 d. Cells cultured on coverslips were fixed and stained in the same way, except for differences in the duration of each operation: fixation (20 min), permeabilization (5 min), washing (3 min), primary and secondary antibody incubation (30 min each).

### Imaging

Images were taken using the following microscopes. For phase-contrast live-imaging, a Leica DMIRE2 inverted microscope equipped with a CCD camera (Sensicam, PCO) controlled by the software package MetaMorph version 7.7.7.0 (Molecular Devices), a stage-top CO2 incubator (Tokai Hit), and an N Plan 2.5×/0.07NA Ph1 lens was used. For imaging of fixed samples, a Nikon A1R inverted confocal microscope controlled by Nikon NIS-Elements software (Nikon), equipped with a Plan Apo 60×/1.40NA lens, was used. For confocal live-imaging, 1) the Nikon A1R microscope equipped with a stage-top CO2 incubator (Tokai Hit) and a Plan Apo VC 20×/0.75NA lens, or 2) an Olympus IX71 inverted microscope equipped with a spinning disk confocal unit (CSU-X1, Yokogawa) and CMOS camera (Orca Flash 4.0, Hamamatsu Photonics) controlled by software package MetaMorph version 7.10.2.240 (Molecular Devices), a stage-top CO2 incubator (Tokai Hit), and a UPlan S Apo 20×/0.75NA lens was used.

### Plasmids and transfection

A vector expressing full-length mouse αE-catenin with a Flag tag at its C-terminus (pCA-αE-catenin–Flag) (67) was a gift from M. Takeichi (RIKEN BDR, Japan). pCA-αE-catenin-Flag (L344P) was constructed using site-specific mutagenesis to introduce a mutation resulting in the single amino acid substitution from L to P (46) into pCA-αE-catenin-Flag. To establish stable clones expressing mutant α-catenin [1–906(L344P)], this vector and PGK-hyg (68) were cotransfected using Lipofectamine 2000 (Invitrogen), and cells were selected by treatment with 400 µg ml^−1^ hygromycin B for 2–4 weeks.

The expression vector of histone H2B tagged with EGFP at the C-terminus was generated using the EGFP-N1 vector (Clontech). The amplified H2B cDNA was cloned into the HindIII/BamHI site of EGFP-N1. To establish cell lines stably expressing GFP-H2B, R2/7 cells stably expressing WT (1-906) or mutant [1–906 (L378P or L344P)] α-catenin were transfected with this vector. EGFP-expressing cells were sorted using a cell sorter (SH800, Sony), sparsely replated, and then single clones were isolated.

### Image analyses

Images were processed and analyzed using ImageJ software (National Institutes of Health) or CellProfiler software (69). To measure spheroid morphology, the outlines of spheroids in phase-contrast images or XY maximum intensity projection images of H2B-GFP (Fig. S7) were traced by freehand drawing, and the circularity (“form factor” in CellProfiler) was calculated using the formula:

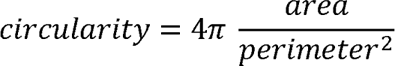

To measure PA length, confocal Z-stack images of α-catenin and F-actin were projected to single images by maximum projection. Spot-like α-catenin-positive objects attached to F-actin bundles were considered PAs. The contour of each PA in the α-catenin channel was traced manually using freehand drawing to define PA regions and max Feret diameters (also known as maximum caliper; the longest distance between two tips along the traced contour of PAs) was measured.

To measure TJ length, confocal Z-stack images of nuclei and ZO-1 were projected as well. The EnhanceOrSuppressFeatures module of CellProfiler enhanced linear structures in the ZO-1channel, and automated segmentation was performed to define the TJ-formed region. The TJ region was then skeletonized to single-pixel width, and the number of pixels representing total TJ length was counted. Cell number was counted using an automated segmentation of nuclei, and TJ length was normalized by dividing using the cell number or field area.

To analyze cell movements during the spheroid formation, 3D DoG (Difference of Gaussian, combined Gaussian radii in sigma of 1.9/3.8 pixels for X/Y dimensions, and 0.7/1.4 pixels for Z dimension) was performed to emphasize nucleus shape and identified distinct nuclei by finding local maxima. These nuclei positions were stored in a SQL database (SQLite3), and their displacements throughout the time series were joined using the nearest neighbor method.

To analyze the morphology of spheroid doublets during fusion, doublet length and contact length measurements were recorded manually by drawing a single line along the long axis of a pair of spheroids or along the fused region (the border of two spheroids), respectively.

### Statistical analyses

Data are expressed as mean ± 95% confidence interval (CI) of at least three independent experiments. All statistical analyses were performed in R software. *P*-values were calculated using Student’s *t*-test for comparison of two groups, and One-way ANOVA followed by Dunnett’s or Tukey’s multiple comparison tests for more than three groups, respectively. In all cases, *P*<0.05 was considered statistically significant.

## Data availability

All data are indicated in the manuscript.

## Supporting information

This article contains supporting information.

## Acknowledgements

We thank Dr. M. Takeichi (RIKEN) for cells and plasmids, Dr. F. Van Roy (Ghent University) for cells, Dr. Sa. Tsukita (Osaka university) for antibodies, and Dr. Y. Hanada (Hirosaki University) for reagents. We thank Drs. K. Horikawa, Y. Hara (Tokushima University), Drs. J. Sakamoto and T. Tomoi (National Institute for Basic Biology) for technical support, K. Aki (Tokushima University) for technical assistance. We also thank Dr. S. Okuda (Kanazawa University) for helpful discussion, Drs. Y. Higuchi (Hirosaki University) and K. Nakagawa (Tokushima University) for helpful comments and encouragement, and all laboratory members for insightful discussion. This work was also supported by Support Center for Advanced Medical Sciences, Tokushima University Graduate School of Biomedical Sciences.

## Author contributions

R. N. and S. Y. conceptualization; R. N. and K. K. data curation; R. N. and K. K. formal analysis; R. N. and S. Y. funding acquisition; R. N., M. S., Y. A., and S. Y. investigation; R. N., K. K., M. S., Y. K., and S. Y. methodology; S. Y. project administration; Y. K., M. T., H. M., and Y. Y resources; K. K. software; S. Y. supervision; R. N. and S. Y. validation; R. N. and K. K. visualization; R. N. writing-original draft; S. Y. writing-review and editing.

## Conflict of Interest

The authors declare that they have no conflicts of interest with the contents of this article.

## Funding and additional information

This work was supported by JSPS KAKENHI (Grant numbers 26650071, 15KT0086, and 18H02617 to S.Y. and 18J22029 to R.N.), Core Research for Evolutional Science and Technology from the Japan Science and Technology Agency (CREST JST; JP115811 to S.Y.), and NIBB Collaborative Research Program (20-518 and 21-422 to K.K. and S.Y.).

## Abbreviations

2D: two-dimensional
3D: three-dimensional
AJ: adherens junction
CE: convergent extension; DAPI, 4’,6-diamidino-2-phenylindole
DMEM: Dulbecco’s modified Eagle medium
ECM: extra cellular matrix
FBS: fetal bovine serum
PA: punctum adherens
PBS: phosphate-buffered saline
PDMS: polydimethylsiloxane
TJ: tight junction
ZA: zonula adherens

## Supplementary figure legends

**Fig. S1. Spheroid formation in V-bottomed microwells and its quantitative analysis. (A)** Still images by time-lapse microscopy. DLD-1 cells (top), R2/7 cells (middle), or R2/7 cells expressing WT α-catenin (1–906; bottom) were seeded on V-bottomed microwells, respectively, and live-imaged for 48 h. Cells expressing WT α-catenin form round spheroids even from initial rectangular shape with high aspect ratio. Scale bar, 250 µm. **(B)** The circularity of spheroid contour was measured and plotted against time. The cadherin-catenin complex function based on α-catenin expression is essential for increase in circularity. Error bars show mean ± 95%CI. **(C)** The circularity of spheroids at 48 h after seeding. Error bars show mean ± 95%CI. (***; *P*<0.001.)

**Fig. S2. Myosin II inhibition does not alter circularity changes of spheroids of both WT and mutant cells. (A)** Still images by time-lapse microscopy. R2/7 cells expressing wild-type (1-906 (WT; top), hypersensitive mutants [1–906 (L378P; middle) or 1–906 (3RE; bottom)] α-catenin were seeded on V-bottomed microwells, respectively, and live-imaged for 48 h in the presence of 50 µM Blebbistatin. Scale bar, 250 µm. **(B)** The circularity of spheroids that measured every 1 h. Error bars show mean ± 95%CI. **(C)** The circularity of spheroids at 48 h after seeding. Error bars show mean ± 95%CI. (***; *P*<0.001.)

**Fig. S3. Inhibition of actin remodeling completely suppresses round spheroid formation. (A)** Still images by time-lapse microscopy. Scale bar, 250 µm. R2/7 cells expressing WT α-catenin were seeded on V-bottomed microwells and live-imaged for 48 h in the presence of 20 µM actin polymerization inhibitor, Cytochalasin D (CytoD; top) or 2 µM actin filament stabilizer, Jasplakinolide (Jasp; bottom), respectively. **(B)** The circularity of spheroids that measured every 1 h. Error bars show mean ± 95%CI. **(C)** The circularity of spheroids at 48 h after seeding. Error bars show mean ± 95%CI. (***; *P*<0.001.)

**Fig. S4. Tension sensitivity mutation of** α**-catenin affects the process of junctional formation after calcium switch. (A)** Visualization of F-actin, FLAG-tagged α-catenin and ZO-1. R2/7 cells expressing wild-type [1–906 (WT; top)], hypersensitive mutant [1–906 (L378P; middle)], or dull mutant [1–906 (L344P; bottom)] α-catenin were seeded on coverslips, respectively, cultured for 24 h in normal medium, and for another 4 h in low calcium medium to dissipate cell-cell adhesion. Then, cells were again cultured in normal medium for 0.5, 1, 2 h, fixed and stained to compare cell-cell adhesion formation process. White arrows indicate PAs. (**B**) Measurement of PA length. Violin plots show data distribution and each dots represents mean values of each biological replicates. **(C)** PA number normalized by area. Each dots represents biological replicates. **(D)** Measurement of TJ length normalized by cell number over time. Each dots represents biological replicates. Scale bars, 5 µm. (**B–D**) Error bars show mean ± 95%CI. (*; *P*<0.05, n.s.; not significant.) See *Materials and Methods* for details of the measurements.

**Fig. S5. Tension sensitivity mutation of** α**-catenin alters junctional formation at cell island peripheries. (A)** Visualization of F-actin, FLAG-tagged α-catenin and ZO-1. R2/7 cells expressing wild-type (1–906 (WT; left), hypersensitive mutant [1–906 (L378P; middle)], or dull mutant [1–906 (L344P; right)] were seeded on coverslips, respectively, cultured for 24 h, fixed, and stained. Scale bars, 10 µm. **(B)** Measurement of PA length. Violin plots show data distribution and each dots represents mean values of each biological replicates. Error bars show mean ± 95%CI. (*; *P*<0.05, ***; *P*<0.001.) See *Materials and Methods* for details of the measurement.

**Fig. S6. Effects of tension sensitivity mutation of α-catenin on the junctional formation of spheroids formed on Matrigel. (A–C)** Visualization of the nucleus, F-actin, FLAG-tagged α-catenin and ZO-1 showing cell-cell junction formation process under 3D culture condition. Images are shown as maximum intensity projection of entire Z-stack except for magnified images (Mag.), which are single slices of the region of the white box. R2/7 cells expressing wild-type [1–906 (WT; **A**)], hypersensitive mutant [1–906 (L378P; **B**)], or dull mutant [1–906 (L344P; **C**)] were seeded on Matrigel, respectively, cultured for 6, 9, 12 h, fixed, and stained. Scale bars, 10 µm.

**Fig. S7. Contribution of folding movement of cell mass to circularity increase. (A)** Still images by confocal time-lapse microscopy (left) and the circularity of spheroids that measured every 10 min (right). Stack images are shown as maximum intensity projection images of the top (XY) or transverse (YZ) view. Time after seeding is indicated in each frame. Yellow arrows indicate the direction of cell mass movement. Scale bar, 100 µm. Each time point indicated in the image is also shown in the graph (black arrows). Red arrows show timings of large increase in the circularity. R2/7 cells expressing H2B-GFP and WT α-catenin were seeded on V-bottomed microwells, respectively, cultured for 6 h and then live-imaged for 11.5 h. Folding movement was often observed in early time (6–12 h) of the rising phase, not in later time (12–17.5 h), consistent with the timing of increase in high correlation movement in **Fig. 7C**. (**B**) Other examples of the circularity changes of spheroids that measured every 10min.

**Fig. S8. Schematic drawing of cellular rearrangement during fusion of two multicellular parts.** Necessity of junctional remodeling (i.e., dissociation and formation) for the fusion of two spheroids is shown. Cell-cell junctions (blue square) between cells numbered 1–3 and 4–6 are not found in the fused resultant spheroid and should have been dissociated, and then, new junctions should be formed between cells 1–6 and 3–4, respectively.

## Supplementary movie legends

**Movie 1. Effects of tension sensitivity mutation of α-catenin on morphogenesis in round-bottomed wells.** Phase-contrast time-lapse imaging of R2/7 cells expressing wild-type [1–906 (WT)] or hypersensitive mutant (1-402) α-catenin seeded in non-adherent round-bottomed wells showing round (WT) or distorted (1-402) morphology. The timestamp represents h:min. Original time interval = 20 min.

**Movie 2. Effects of tension sensitivity mutation of** α**-catenin on morphogenesis in round-bottomed wells.** Phase-contrast time-lapse imaging of R2/7 cells expressing wild-type [1–906 (WT)] or hypersensitive mutant [1–906 (R326E or R548E or R551E or 3RE or M319G or L378P or M 319G/L378P)] α-catenin seeded in non-adherent round-bottomed wells showing different degree of distortion according to tension sensitivity. The timestamp represents h:min. Original time interval = 1 h.

**Movie 3. Initial aspect ratios of cell mass and final spheroid circularity.** Phase-contrast time-lapse imaging of DLD-1 cells seeded in V-bottomed microwells with different aspect ratios. Aspect ratios are 1:1, 1.5:1, 2:1 or 3:1from top, respectively. Final shape of spheroids and timings of spheroid rounding was constant among all aspect ratios. The timestamp represents h:min. original time interval = 10 min.

**Movie 4. Effects of α-catenin depletion on morphogenesis in V-bottomed microwells.** Phase-contrast time-lapse imaging of DLD-1 cells, R2/7 cells, and R2/7 cells expressing wild-type [1-906 (WT)] α-catenin seeded on V-bottomed microwells. The timestamp represents h:min. Original time interval = 1 min.

**Movie 5. Effects of tension sensitivity increase of α-catenin on morphogenesis in V-bottomed microwells.** Phase-contrast time-lapse imaging of R2/7 cells expressing hypersensitive mutant α-catenin [1–906 (R326E/R548E/R551E; 3RE), 1–402, 1–906 (L378P)] seeded on V-bottomed microwells. The timestamp represents h:min. Original time interval = 1 min.

**Movie 6. Effects of tension sensitivity decrease of α-catenin on morphogenesis in V-bottomed microwells.** Phase-contrast time-lapse imaging of R2/7 cells expressing dull mutant α-catenin [1–906 (L344P)] seeded on V-bottomed microwells. The timestamp represents h:min. Original time interval = 1 min.

**Movie 7. Effects of myosin II inhibition on morphogenesis in V-bottomed microwells.** Phase-contrast time-lapse imaging of R2/7 cells expressing wild-type [1–906 (WT)], or hypersensitive mutant [1–906 (L378P) or 1–906 (3RE)] α-catenin in the presence of 50 µM Blebbistatin, respectively. The timestamp represents h:min. Original time interval = 1 min.

**Movie 8. Effects of actin dynamics inhibition on morphogenesis in V-bottomed microwells.** Phase-contrast time-lapse imaging of R2/7 cells expressing wild-type [1–906 (WT)] α-catenin in the presence of 20 µM Cytochalasin D, an inhibitor of actin polymerization or 2 µM Jasplakinolide, an actin filament stabilizer. The timestamp represents h:min. Original time interval = 1 min.

**Movie 9. Effects of Dynamin inhibition on morphogenesis in V-bottomed microwells.** Phase-contrast time-lapse imaging of R2/7 cells expressing wild-type [1–906 (WT)], hypersensitive mutant [1–906 (L378P)], or dull mutant [1–906(L344P)] α-catenin in the presence of 80 µM Dynasore, respectively. The timestamp represents h:min. Original time interval = 1 min.

**Movie 10. Visualization of cellular tracking during spheroid formation.** The positions of nucleus and displacement correlations of neighboring cell pairs detected from spinning-disk confocal time-lapse images are visualized as follows. Each dot indicates nucleus position in the XY plane, and its depth is expressed as a color of each dot (‘colder’ color shows deeper (closer to bottom) position). Displacement correlation of each neighboring cell pair (connected by lines) is also shown by the same color scale (‘colder’ color shows negative, ‘hotter’ color shows a positive correlation, respectively). Frame numbers are indicated at top right. Time interval = 10 sec.

**Movie 11. Effects of tension sensitivity mutation of α-catenin or Dynamin inhibition on cell behavior during spheroid formation.** Spinning-disk confocal time-lapse imaging in GFP-H2B to visualize nucleus in wild-type [1–906 (WT)], hypersensitive mutant [1–906 (L378P)], or dull mutant [1–906(L344P)] α-catenin expressing R2/7 cells. Wild-type α-catenin expressing R2/7 cells cultured in the presence of 10 µM MiTMAB is also shown. The movie shows a maximum intensity projection in XY and ZY plane of 42 optical slices acquired at a 3.6 µm step size. The timestamp represents h:min:sec. Original time interval = 10 sec.

**Movie 12. Effects of tension sensitivity mutation of α-catenin on spheroid fusion.** Confocal bright-field time-lapse imaging in R2/7 cells expressing wild-type [1–906 (WT)], hypersensitive mutant [1–906 (L378P)], or dull mutant [1–906(L344P)] α-catenin. The movie shows a single slice. The timestamp represents h:min. Time interval = 5 min.

## Notes

### Competing Interest Statement

The authors have declared no competing interest.

### Summary of Updates

Conversion to a PDF file was not successful for Fig.7. We corrected this.

## References

1. Honda, H. (2017) The world of epithelial sheets. Dev Growth Differ. 59, 306–316

2. Herbst, C. (1900) über das Auseinandergehen von Furchungs- und Gewebezellen in kalkfreiem Medium. Archiv Für Entwicklungsmechanik Der Org. 9, 424–463

3. Wilson, H. V. (1907) On some phenomena of coalescence and regeneration in sponges. J Exp Zool. 5, 245–258

4. Steinberg, M. S. (1963) Reconstruction of Tissues by Dissociated Cells. Science. 141, 401–408

5. Harris, A. K. (1976) Is cell sorting caused by differences in the work of intercellular adhesion? A critique of the steinberg hypothesis. J Theor Biol. 61, 267–285

6. Krieg, M., Arboleda-Estudillo, Y., Puech, P.-H., Käfer, J., Graner, F., Müller, D. J., and Heisenberg, C.-P. (2008) Tensile forces govern germ-layer organization in zebrafish. Nat Cell Biol. 10, 429–436

7. Winklbauer, R. (2015) Cell adhesion strength from cortical tension – an integration of concepts. J Cell Sci. 128, 3687–3693

8. Voss, A. K., and Strasser, A. (2020) The essentials of developmental apoptosis. F1000research. 9, F1000 Faculty Rev-148

9. Godard, B. G., and Heisenberg, C.-P. (2019) Cell division and tissue mechanics. Curr Opin Cell Biol. 60, 114–120

10. Scarpa, E., and Mayor, R. (2016) Collective cell migration in development. J Cell Biol. 212, 143–155

11. Paluch, E., and Heisenberg, C.-P. (2009) Biology and Physics of Cell Shape Changes in Development. Curr Biol. 19, R790–R799

12. Basson, M. A. (2012) Signaling in Cell Differentiation and Morphogenesis. Csh Perspect Biol. 4, a008151

13. Quintin, S., Gally, C., and Labouesse, M. (2008) Epithelial morphogenesis in embryos: asymmetries, motors and brakes. Trends Genet. 24, 221–230

14. Wang, Y.-C. (2021) The origin and the mechanism of mechanical polarity during epithelial folding. Semin Cell Dev Biol. 10.1016/j.semcdb.2021.05.027

15. Glazier, J. A., and Graner, F. (1993) Simulation of the differential adhesion driven rearrangement of biological cells. Phys Rev E. 47, 2128–2154

16. Hirashima, T., Rens, E. G., and Merks, R. M. H. (2017) Cellular Potts modeling of complex multicellular behaviors in tissue morphogenesis. Dev Growth Differ. 59, 329–339

17. Honda, H., and Nagai, T. (2014) Cell models lead to understanding of multi-cellular morphogenesis consisting of successive self-construction of cells. J Biochem. 157, 129–136

18. Ishihara, S., Marcq, P., and Sugimura, K. (2017) From cells to tissue: A continuum model of epithelial mechanics. Phys Rev E. 96, 022418

19. Okuda, S., Inoue, Y., and Adachi, T. (2015) Three-dimensional vertex model for simulating multicellular morphogenesis. Biophysics Physicobiology. 12, 13–20

20. Yonemura, S. (2017) Actin filament association at adherens junctions. J Medical Investigation. 64, 14–19

21. Lecuit, T., and Yap, A. S. (2015) E-cadherin junctions as active mechanical integrators in tissue dynamics. Nat Cell Biol. 17, 533 539

22. Takeichi, M. (2014) Dynamic contacts: rearranging adherens junctions to drive epithelial remodelling. Nat Rev Mol Cell Bio. 15, 397 410

23. Martin, A. C., and Goldstein, B. (2014) Apical constriction: themes and variations on a cellular mechanism driving morphogenesis. Development. 141, 1987–1998

24. Vasquez, C. G., and Martin, A. C. (2016) Force transmission in epithelial tissues. Dev Dynam. 245, 361–371

25. Pokutta, S., and Weis, W. I. (2007) Structure and Mechanism of Cadherins and Catenins in Cell-Cell Contacts. Cell Dev Biology. 23, 237–261

26. Takeichi, M. (1995) Morphogenetic roles of classic cadherins. Curr Opin Cell Biol. 7, 619–627

27. Desai, R., Sarpal, R., Ishiyama, N., Pellikka, M., Ikura, M., and Tepass, U. (2013) Monomeric α-catenin links cadherin to the actin cytoskeleton. Nat Cell Biol. 15, 261–273

28. Ishiyama, N., Sarpal, R., Wood, M. N., Barrick, S. K., Nishikawa, T., Hayashi, H., Kobb, A. B., Flozak, A. S., Yemelyanov, A., Fernandez-Gonzalez, R., Yonemura, S., Leckband, D. E., Gottardi, C. J., Tepass, U., and Ikura, M. (2018) Force-dependent allostery of the α-catenin actin-binding domain controls adherens junction dynamics and functions. Nat Commun. 9, 5121

29. Yonemura, S., Wada, Y., Watanabe, T., Nagafuchi, A., and Shibata, M. (2010) α-Catenin as a tension transducer that induces adherens junction development. Nat Cell Biol. 12, 533 542

30. Miyake, Y., Inoue, N., Nishimura, K., Kinoshita, N., Hosoya, H., and Yonemura, S. (2006) Actomyosin tension is required for correct recruitment of adherens junction components and zonula occludens formation. Exp Cell Res. 312, 1637 1650

31. Duc, Q. le, Shi, Q., Blonk, I., Sonnenberg, A., Wang, N., Leckband, D., and Rooij, J. de (2010) Vinculin potentiates E-cadherin mechanosensing and is recruited to actin-anchored sites within adherens junctions in a myosin II–dependent manner. J Cell Biol. 189, 1107–1115

32. Hirano, S., Kimoto, N., Shimoyama, Y., Hirohashi, S., and Takeichi, M. (1992) Identification of a neural α-catenin as a key regulator of cadherin function and multicellular organization. Cell. 70, 293–301

33. Torres, M., Stoykova, A., Huber, O., Chowdhury, K., Bonaldo, P., Mansouri, A., Butz, S., Kemler, R., and Gruss, P. (1997) An α-E-catenin gene trap mutation defines its function in preimplantationLdevelopment. Proc National Acad Sci. 94, 901–906

34. Vasioukhin, V., Bauer, C., Degenstein, L., Wise, B., and Fuchs, E. (2001) Hyperproliferation and Defects in Epithelial Polarity upon Conditional Ablation of α-Catenin in Skin. Cell. 104, 605–617

35. Watabe, M., Nagafuchi, A., Tsukita, S., and Takeichi, M. (1994) Induction of polarized cell-cell association and retardation of growth by activation of the E-cadherin-catenin adhesion system in a dispersed carcinoma line. J Cell Biology. 127, 247–256

36. Watabe-Uchida, M., Uchida, N., Imamura, Y., Nagafuchi, A., Fujimoto, K., Uemura, T., Vermeulen, S., Roy, F. van, Adamson, E. D., and Takeichi, M. (1998) α-Catenin-Vinculin Interaction Functions to Organize the Apical Junctional Complex in Epithelial Cells. J Cell Biol. 142, 847–857

37. Matsuzawa, K., Himoto, T., Mochizuki, Y., and Ikenouchi, J. (2018) α-Catenin Controls the Anisotropy of Force Distribution at Cell-Cell Junctions during Collective Cell Migration. Cell Reports. 23, 3447–3456

38. Sakakibara, S., Mizutani, K., Sugiura, A., Sakane, A., Sasaki, T., Yonemura, S., and Takai, Y. (2020) Afadin regulates actomyosin organization through αE-catenin at adherens junctions. J Cell Biol. 10.1083/jcb.201907079

39. Itoh, M., Nagafuchi, A., Moroi, S., and Tsukita, S. (1997) Involvement of ZO-1 in Cadherin-based Cell Adhesion through Its Direct Binding to α Catenin and Actin Filaments. J Cell Biology. 138, 181–192

40. Xu, W., Baribault, H., and Adamson, E. D. (1998) Vinculin knockout results in heart and brain defects during embryonic development. Dev Camb Engl. 125, 327–37

41. Hirano, Y., Amano, Y., Yonemura, S., and Hakoshima, T. (2018) The forceLsensing device region of αLcatenin is an intrinsically disordered segment in the absence of intramolecular stabilization of the autoinhibitory form. Genes Cells. 23, 370–385

42. Rangarajan, E. S., and Izard, T. (2013) Dimer asymmetry defines α-catenin interactions. Nat Struct Mol Biol. 20, 188 193

43. Yao, M., Qiu, W., Liu, R., Efremov, A. K., Cong, P., Seddiki, R., Payre, M., Lim, C. T., Ladoux, B., Mège, R.-M., and Yan, J. (2014) Force-dependent conformational switch of α-catenin controls vinculin binding. Nat Commun. 5, 4525

44. Maki, K., Han, S.-W., Hirano, Y., Yonemura, S., Hakoshima, T., and Adachi, T. (2018) Real-time TIRF observation of vinculin recruitment to stretched α-catenin by AFM. Sci Rep-uk. 8, 1575

45. Li, J., Newhall, J., Ishiyama, N., Gottardi, C., Ikura, M., Leckband, D. E., and Tajkhorshid, E. (2015) Structural Determinants of the Mechanical Stability of α-Catenin. J Biol Chem. 290, 18890–18903

46. Peng, X., Maiers, J. L., Choudhury, D., Craig, S. W., and DeMali, K. A. (2012) α-Catenin Uses a Novel Mechanism to Activate Vinculin. J Biol Chem. 287, 7728–7737

47. Seddiki, R., Narayana, G. H. N. S., Strale, P.-O., Balcioglu, H. E., Peyret, G., Yao, M., Le, A. P., Teck, L. C., Yan, J., Ladoux, B., and Mège, R. M. (2018) Force-dependent binding of vinculin to α-catenin regulates cell-cell contacts stability and collective cell behavior. Mol Biol Cell. 29, mbc.E17-04-0231

48. Tada, M., and Heisenberg, C.-P. (2012) Convergent extension: using collective cell migration and cell intercalation to shape embryos. Development. 139, 3897–3904

49. Han, M. K. L., Hoijman, E., Nöel, E., Garric, L., Bakkers, J., and Rooij, J. de (2016) αE-catenin-dependent mechanotransduction is essential for proper convergent extension in zebrafish. Biol Open. 5, bio.021378

50. Hengel, J. van, Gohon, L., Bruyneel, E., Vermeulen, S., Cornelissen, M., Mareel, M., and Roy, F. van (1997) Protein Kinase C Activation Upregulates Intercellular Adhesion of α-Catenin–negative Human Colon Cancer Cell Variants via Induction of Desmosomes. J Cell Biol. 137, 1103–1116

51. Carlier, M.-F., and Shekhar, S. (2017) Global treadmilling coordinates actin turnover and controls the size of actin networks. Nat Rev Mol Cell Bio. 18, 389–401

52. Yonemura, S., Itoh, M., Nagafuchi, A., and Tsukita, S. (1995) Cell-to-cell adherens junction formation and actin filament organization: similarities and differences between non-polarized fibroblasts and polarized epithelial cells. J Cell Sci. 108 ( **Pt 1**), 127–42

53. Hong, S., Troyanovsky, R. B., and Troyanovsky, S. M. (2010) Spontaneous assembly and active disassembly balance adherens junction homeostasis. Proc National Acad Sci. 107, 3528–3533

54. Kametani, Y., and Takeichi, M. (2007) Basal-to-apical cadherin flow at cell junctions. Nat Cell Biol. 9, 92–98

55. Levayer, R., Pelissier-Monier, A., and Lecuit, T. (2011) Spatial regulation of Dia and Myosin-II by RhoGEF2 controls initiation of E-cadherin endocytosis during epithelial morphogenesis. Nat Cell Biol. 13, 529–40

56. Beco, S. de, Perney, J.-B., Coscoy, S., and Amblard, F. (2015) Mechanosensitive Adaptation of E-Cadherin Turnover across adherens Junctions. Plos One. 10, e0128281

57. Iyer, K. V., Piscitello-Gómez, R., Paijmans, J., Jülicher, F., and Eaton, S. (2019) Epithelial Viscoelasticity Is Regulated by Mechanosensitive E-cadherin Turnover. Current Biology. 29, 578–591.e5

58. Zhu, M., and Zernicka-Goetz, M. (2020) Principles of Self-Organization of the Mammalian Embryo. Cell. 183, 1467–1478

59. Lecuit, T., and Lenne, P.-F. (2007) Cell surface mechanics and the control of cell shape, tissue patterns and morphogenesis. Nat Rev Mol Cell Bio. 8, 633–644

60. Cavo, M., Cave, D. D., D’Amone, E., Gigli, G., Lonardo, E., and Mercato, L. L. del (2020) A synergic approach to enhance long-term culture and manipulation of MiaPaCa-2 pancreatic cancer spheroids. Sci Rep-uk. 10, 10192

61. Deckers, T., Lambrechts, T., Viazzi, S., Hall, G. N., Papantoniou, I., Bloemen, V., and Aerts, J.-M. (2018) High-throughput image-based monitoring of cell aggregation and microspheroid formation. Plos One. 13, e0199092

62. Raghavan, S., Mehta, P., Horst, E. N., Ward, M. R., Rowley, K. R., and Mehta, G. (2016) Comparative analysis of tumor spheroid generation techniques for differential in vitro drug toxicity. Oncotarget. 7, 16948–16961

63. Smeets, B., Pešek, J., Deckers, T., Hall, G. N., Cuvelier, M., Ongenae, S., Bloemen, V., Luyten, F. P., Papantoniou, I., and Ramon, H. (2020) Compaction Dynamics during Progenitor Cell Self-Assembly Reveal Granular Mechanics. Matter. 2, 1283–1295

64. Maki, K., Han, S.-W., Hirano, Y., Yonemura, S., Hakoshima, T., and Adachi, T. (2016) Mechano-adaptive sensory mechanism of α-catenin under tension. Sci Rep-uk. 6, 24878

65. Wu, Z., and Hjort, K. (2009) Surface modification of PDMS by gradient-induced migration of embedded Pluronic. Lab Chip. 9, 1500–1503

66. Wang, J.-C., Liu, W., Tu, Q., Ma, C., Zhao, L., Wang, Y., Ouyang, J., Pang, L., and Wang, J. (2014) High throughput and multiplex localization of proteins and cells for in situ micropatterning using pneumatic microfluidics. Analyst. 140, 827–836

67. Abe, K., Chisaka, O., Roy, F. van, and Takeichi, M. (2004) Stability of dendritic spines and synaptic contacts is controlled by αN-catenin. Nat Neurosci. 7, 357–363

68. Mortensen, R. M., Zubiaur, M., Neer, E. J., and Seidman, J. G. (1991) Embryonic stem cells lacking a functional inhibitory G-protein subunit (alpha i2) produced by gene targeting of both alleles. Proc National Acad Sci. 88, 7036–7040

69. Carpenter, A. E., Jones, T. R., Lamprecht, M. R., Clarke, C., Kang, I. H., Friman, O., Guertin, D. A., Chang, J. H., Lindquist, R. A., Moffat, J., Golland, P., and Sabatini, D. M. (2006) CellProfiler: image analysis software for identifying and quantifying cell phenotypes. Genome Biol. 7, R100

