## Supporting Information for "Appropriate tension sensitivity of α-catenin ensures rounding morphogenesis of epithelial spheroids"

### Supplementary figures

Fig. S1. Spheroid formation in V-bottomed microwells and its quantitative analysis.

Fig. S2. Myosin II inhibition does not alter circularity changes of spheroids of both WT and mutant cells.

Fig. S3. Inhibition of actin remodeling completely suppresses round spheroid formation.

Fig. S4. Tension sensitivity mutation of α-catenin affects the process of junctional formation after calcium switch.

Fig. S5. Tension sensitivity mutation of α-catenin alters junctional formation at cell island peripheries.

Fig. S6. Effects of tension sensitivity mutation of α-catenin on the junctional formation of spheroids formed on Matrigel.

Fig. S7. Contribution of folding movement of cell mass to circularity increase.

Fig. S8. Schematic drawing of cellular rearrangement during fusion of two multicellular parts.

### Supplementary movies

Movie 1. Effects of tension sensitivity mutation of α-catenin on morphogenesis in round-bottomed wells.

Movie 2. Effects of tension sensitivity mutation of α-catenin on morphogenesis in round-bottomed wells.

Movie 3. Initial aspect ratios of cell mass and final spheroid circularity.

Movie 4. Effects of α-catenin depletion on morphogenesis in V-bottomed microwells.

Movie 5. Effects of tension sensitivity increase of α-catenin on morphogenesis in V-bottomed microwells.

Movie 6. Effects of tension sensitivity decrease of α-catenin on morphogenesis in V-bottomed microwells.

Movie 7. Effects of myosin II inhibition on morphogenesis in V-bottomed microwells.

Movie 8. Effects of actin dynamics inhibition on morphogenesis in V-bottomed microwells.

Movie 9. Effects of Dynamin inhibition on morphogenesis in V-bottomed microwells.

Movie 10. Visualization of cellular tracking during spheroid formation.

Movie 11. Effects of tension sensitivity mutation of α-catenin or Dynamin inhibition on cell behavior during spheroid formation.

Movie 12. Effects of tension sensitivity mutation of α-catenin on spheroid fusion.

### Supplementary figures

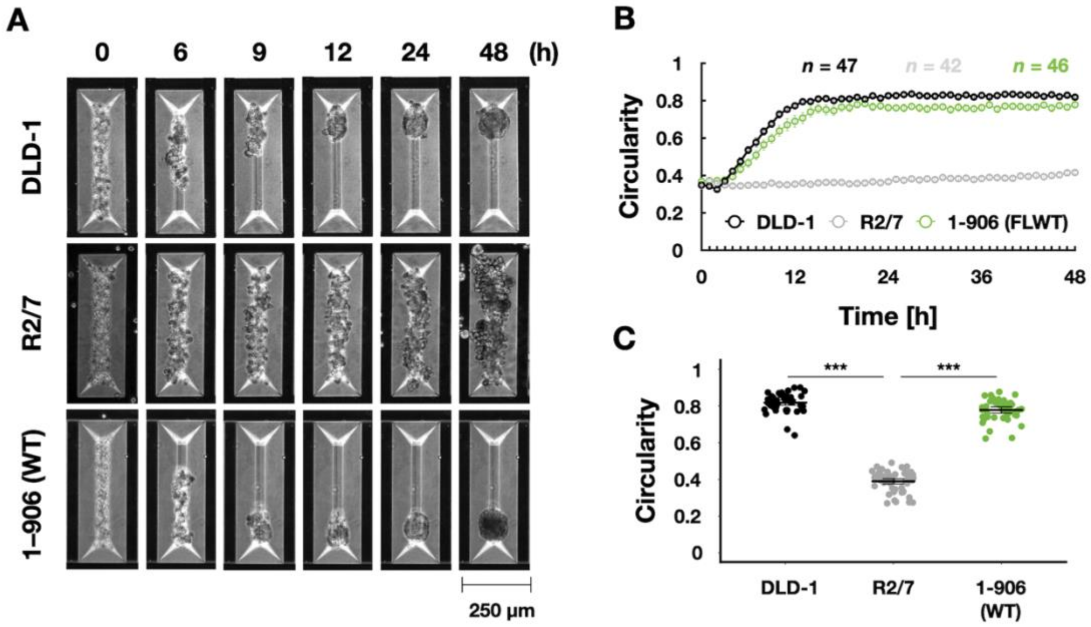


**Fig. S1. Spheroid formation in V-bottomed microwells and its quantitative analysis.**

**(A)** Still images by time-lapse microscopy. DLD-1 cells (top), R2/7 cells (middle), or R2/7 cells expressing WT α-catenin (1–906; bottom) were seeded on V-bottomed microwells, respectively, and live-imaged for 48 h. Cells expressing WT α-catenin form round spheroids even from initial rectangular shape with high aspect ratio. Scale bar, 250 µm. **(B)** The circularity of spheroid contour was measured and plotted against time. The cadherin-catenin complex function based on α-catenin expression is essential for increase in circularity. Error bars show mean ± 95%CI. **(C)** The circularity of spheroids at 48 h after seeding. Error bars show mean ± 95%CI. (***; *P*<0.001.)

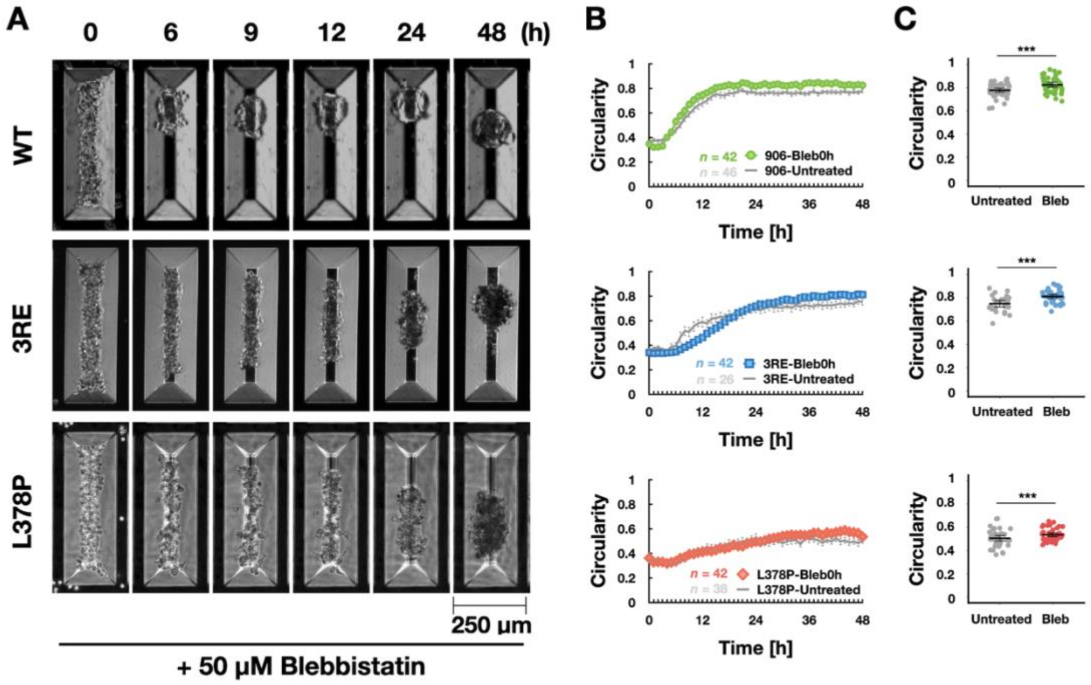


**Fig. S2. Myosin II inhibition does not alter circularity changes of spheroids of both WT and mutant cells.**

**(A)** Still images by time-lapse microscopy. R2/7 cells expressing wild-type (1-906 (WT; top), hypersensitive mutants [1–906 (L378P; middle) or 1–906 (3RE; bottom)] α-catenin were seeded on V-bottomed microwells, respectively, and live-imaged for 48 h in the presence of 50 μM Blebbistatin. Scale bar, 250 μm. **(B)** The circularity of spheroids that measured every 1 h. Error bars show mean ± 95%CI. **(C)** The circularity of spheroids at 48 h after seeding. Error bars show mean ± 95%CI. (***; $P<0.001$.)

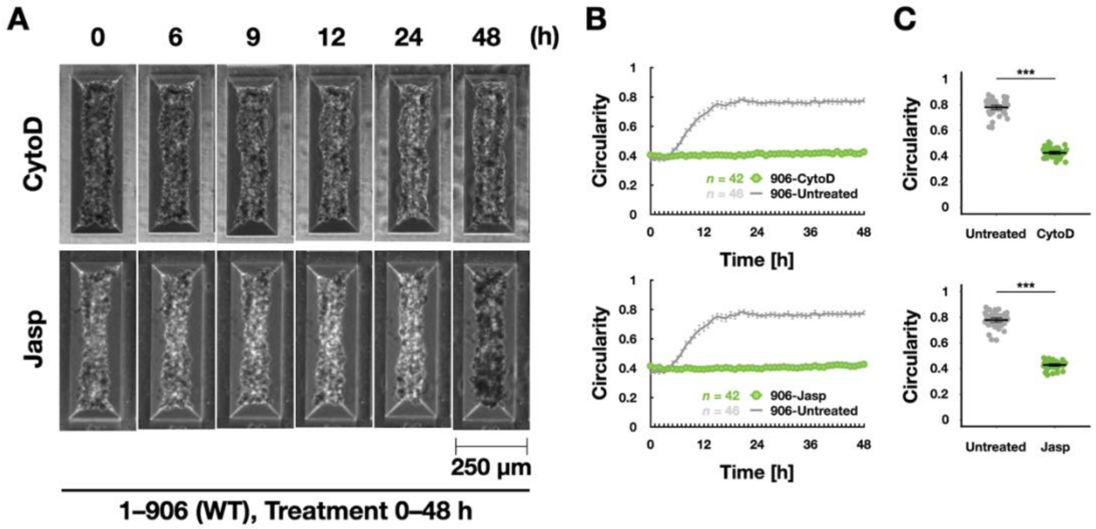


**Fig. S3. Inhibition of actin remodeling completely suppresses round spheroid formation.**

**(A)** Still images by time-lapse microscopy. Scale bar, 250 µm. R2/7 cells expressing WT α-catenin were seeded on V-bottomed microwells and live-imaged for 48 h in the presence of 20 µM actin polymerization inhibitor, Cytochalasin D (CytoD; top) or 2 µM actin filament stabilizer, Jasplakinolide (Jasp; bottom), respectively. **(B)** The circularity of spheroids that measured every 1 h. Error bars show mean ± 95%CI. **(C)** The circularity of spheroids at 48 h after seeding. Error bars show mean ± 95%CI. (***; $P<0.001$.)

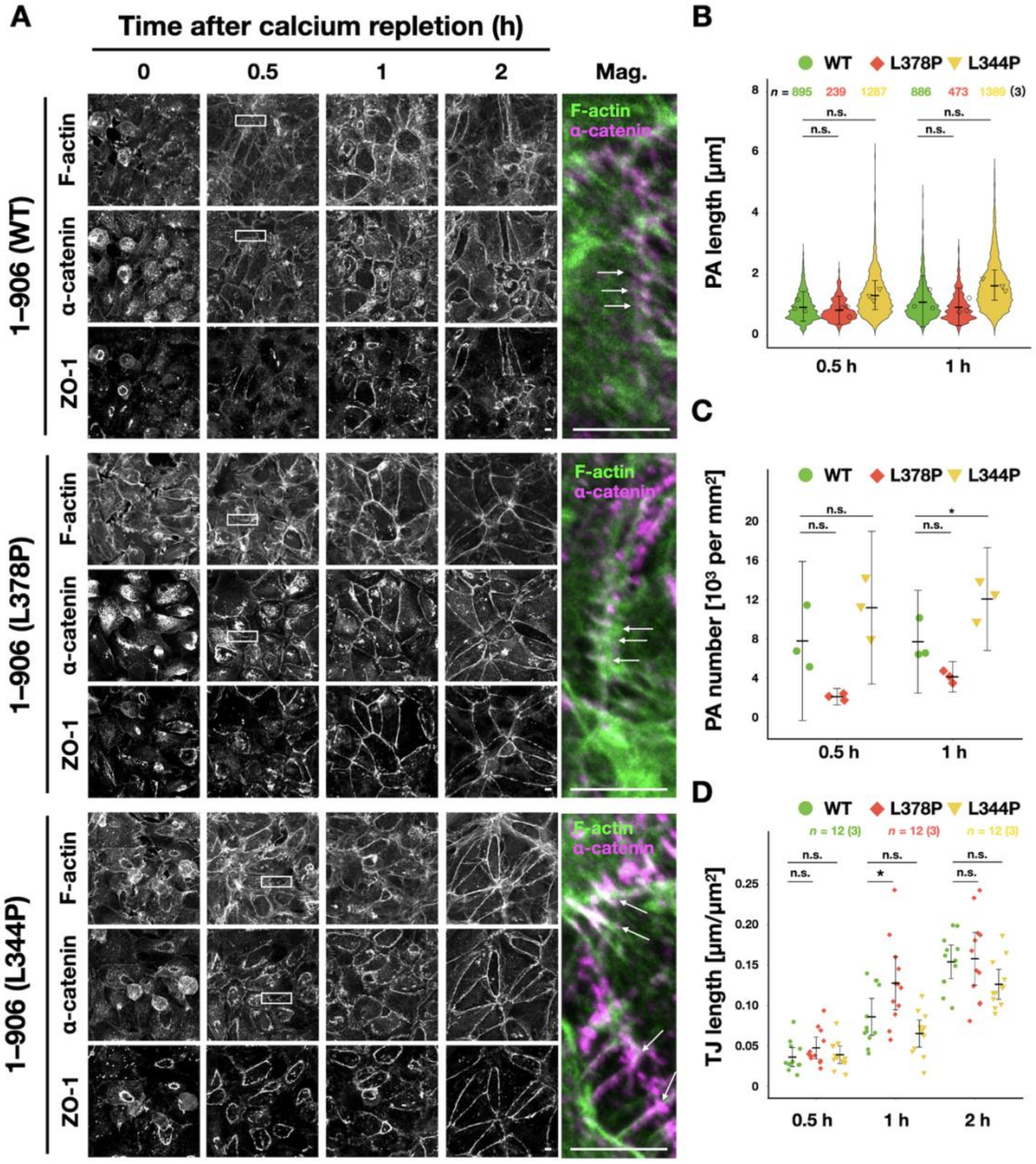


**Fig. S4. Tension sensitivity mutation of α-catenin affects the process of junctional formation after calcium switch.**

**(A)** Visualization of F-actin, FLAG-tagged α-catenin and ZO-1. R2/7 cells expressing wild-type [1–906 (WT; top)], hypersensitive mutant [1–906 (L378P; middle)], or dull mutant [1–906 (L344P; bottom)] α-catenin were seeded on coverslips, respectively, cultured for 24 h in normal medium, and for another 4 h in low calcium medium to dissipate cell-cell adhesion. Then, cells were again cultured in normal medium for 0.5, 1, 2 h, fixed and stained to compare cell-cell adhesion formation process. White arrows indicate PAs. (**B**) Measurement of PA length. Violin plots show data distribution and each dots represents mean values of each biological replicates. **(C)** PA number normalized by area. Each dots represents biological replicates. **(D)** Measurement of TJ length normalized by cell number over time**.** Each dots represents biological replicates. Scale bars, 5 μm. (**B–D**) Error bars show mean ± 95%CI. (*; $P<0.05$, n.s.; not significant.) See *Materials and Methods* for details of the measurements.

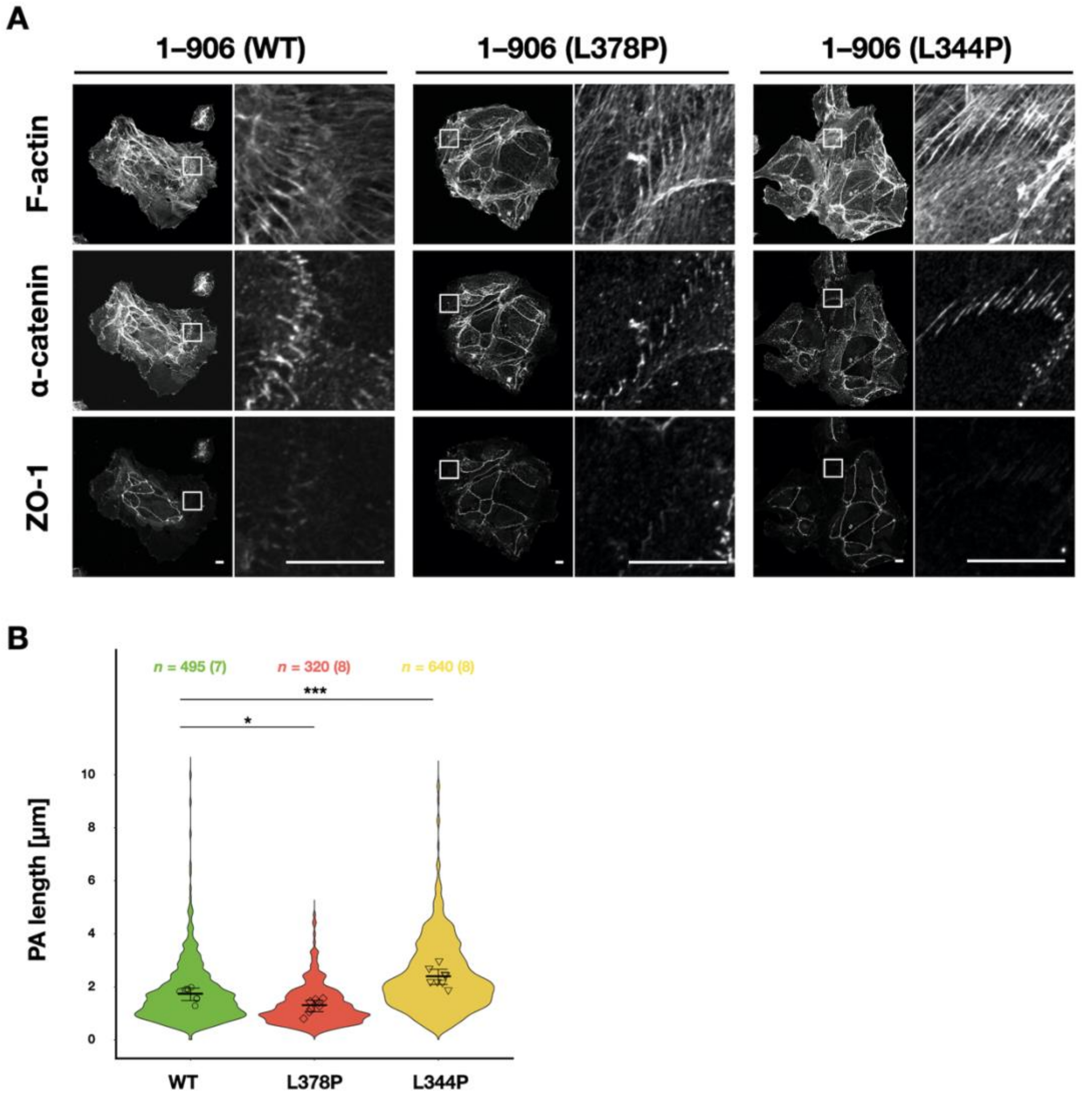


**Fig. S5. Tension sensitivity mutation of α-catenin alters junctional formation at cell island peripheries.**

**(A)** Visualization of F-actin, FLAG-tagged α-catenin and ZO-1. R2/7 cells expressing wild-type (1–906 (WT; left), hypersensitive mutant [1–906 (L378P; middle)], or dull mutant [1–906 (L344P; right)] were seeded on coverslips, respectively, cultured for 24 h, fixed, and stained. Scale bars, 10 µm. **(B)** Measurement of PA length. Violin plots show data distribution and each dots represents mean values of each biological replicates. Error bars show mean ± 95%CI. (*; *P*<0.05, ***; *P*<0.001.) See *Materials and Methods* for details of the measurement.

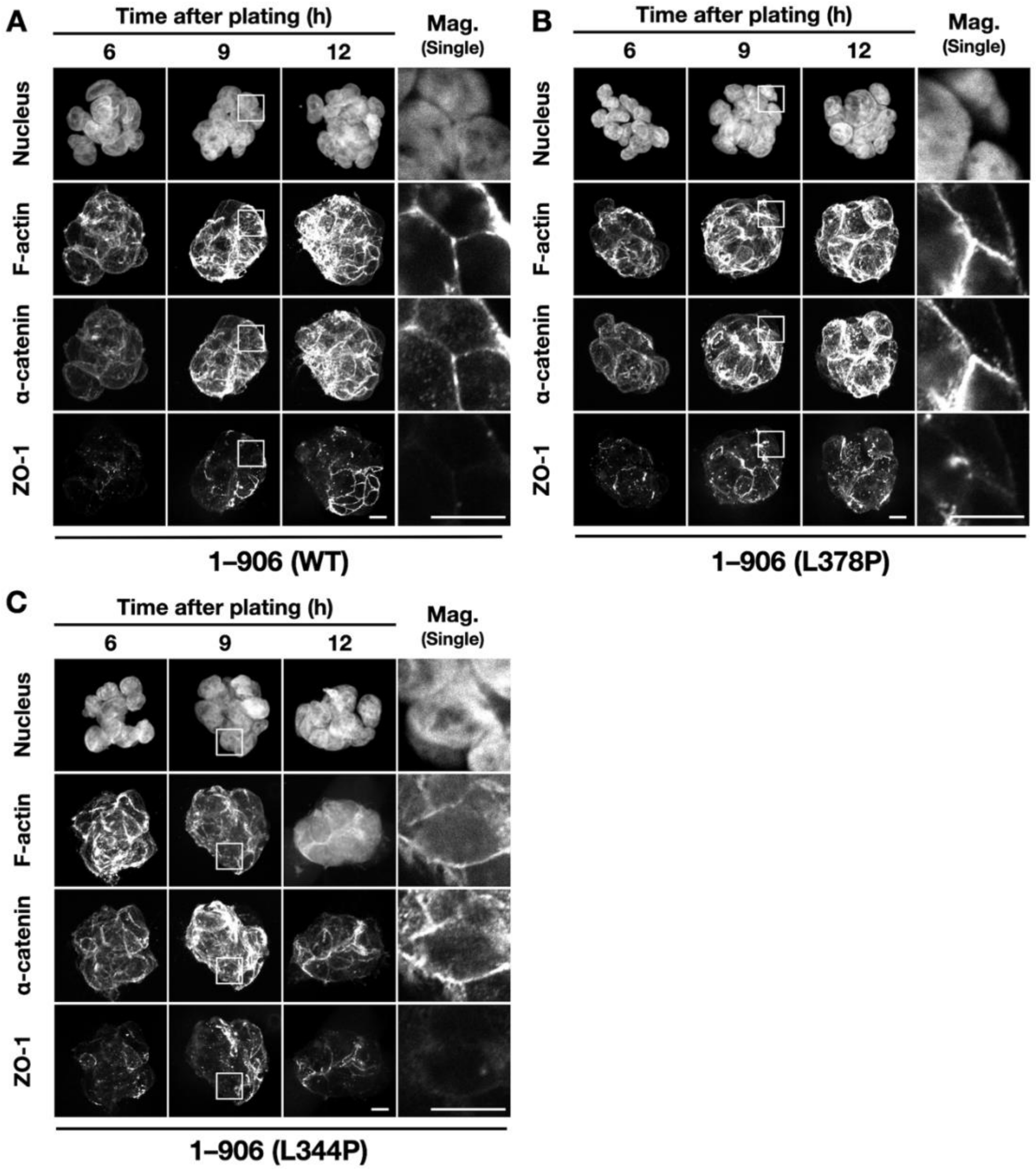


**Fig. S6. Effects of tension sensitivity mutation of α-catenin on the junctional formation of spheroids formed on Matrigel.**

**(A–C)** Visualization of the nucleus, F-actin, FLAG-tagged α-catenin and ZO-1 showing cell-cell junction formation process under 3D culture condition. Images are shown as maximum intensity projection of entire Z-stack except for magnified images (Mag.), which are single slices of the region of the white box. R2/7 cells expressing wild-type [1–906 (WT; **A**)], hypersensitive mutant [1–906 (L378P; **B**)], or dull mutant [1–906 (L344P; **C**)] were seeded on Matrigel, respectively, cultured for 6, 9, 12 h, fixed, and stained. Scale bars, 10 μm.

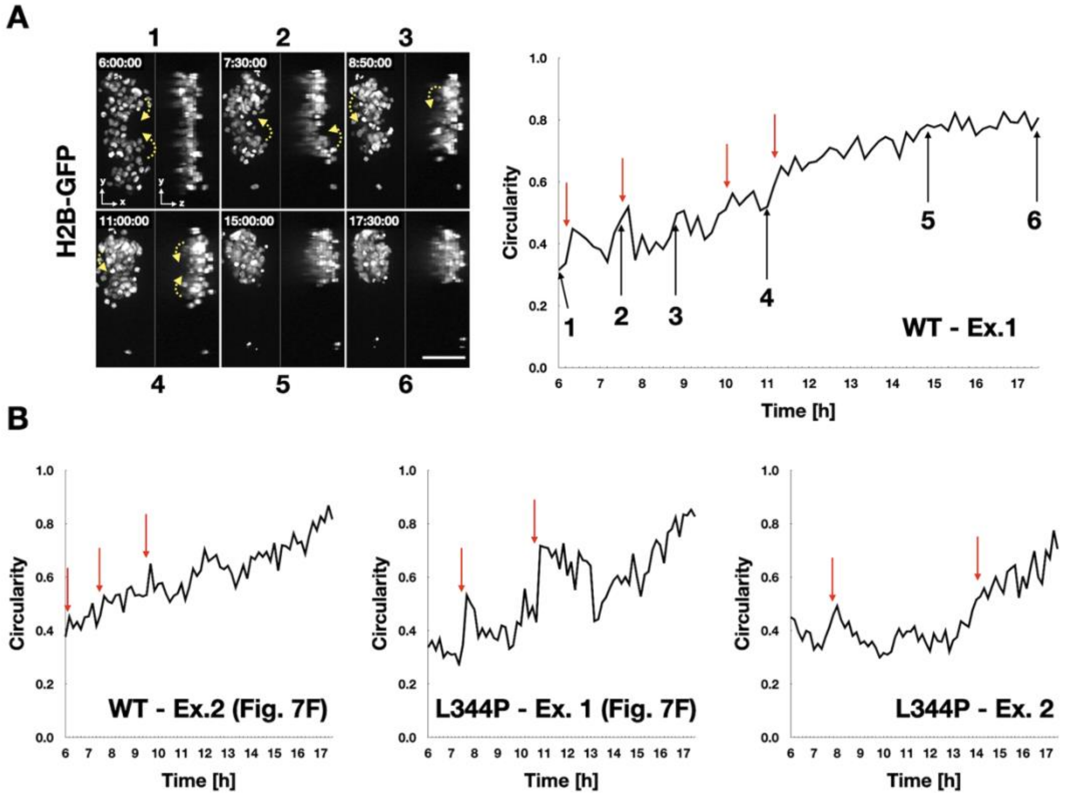


**Fig. S7. Contribution of folding movement of cell mass to circularity increase.**

**(A)** Still images by confocal time-lapse microscopy (left) and the circularity of spheroids that measured every 10 min (right). Stack images are shown as maximum intensity projection images of the top (XY) or transverse (YZ) view. Time after seeding is indicated in each frame. Yellow arrows indicate the direction of cell mass movement. Scale bar, 100 µm. Each time point indicated in the image is also shown in the graph (black arrows). Red arrows show timings of large increase in the circularity. R2/7 cells expressing H2B-GFP and WT α-catenin were seeded on V-bottomed microwells, respectively, cultured for 6 h and then live-imaged for 11.5 h. Folding movement was often observed in early time (6–12 h) of the rising phase, not in later time (12–17.5 h), consistent with the timing of increase in high correlation movement in **Fig. 7C**. (**B**) Other examples of the circularity changes of spheroids that measured every 10min.

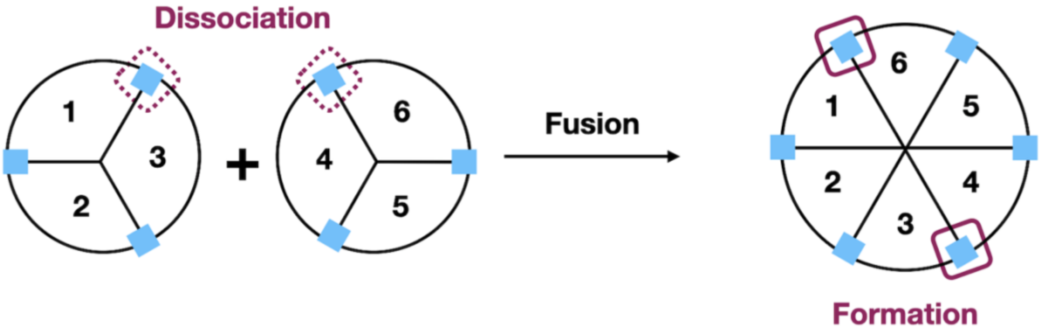


**Fig. S8. Schematic drawing of cellular rearrangement during fusion of two multicellular parts.**

Necessity of junctional remodeling (i.e., dissociation and formation) for the fusion of two spheroids is shown. Cell-cell junctions (blue square) between cells numbered 1–3 and 4–6 are not found in the fused resultant spheroid and should have been dissociated, and then, new junctions should be formed between cells 1–6 and 3–4, respectively.
